# Critical Mechanistic Features of HIV-1 Viral Capsid Assembly

**DOI:** 10.1101/2022.05.03.490470

**Authors:** Manish Gupta, Alexander J. Pak, Gregory A. Voth

**Author notes:** Department of Chemical and Biological Engineering, Colorado School of Mines, Golden, CO 80401, United States.

## Abstract

The maturation of HIV-1 capsid protein (CA) into a cone-shaped lattice encasing the condensed RNA genome is critical for viral infectivity. During HIV-1 maturation, CA can self-assemble into a wide range of capsid morphologies made of ~175-250 hexamers and 12 pentamers. Most recently, the cellular polyanion inositol hexakisphosphate (IP6) has been demonstrated to facilitate conical capsid formation by coordinating a ring of arginine residues within the central cavity of capsid hexamers and pentamers. However, the precise kinetic interplay of events during IP6 and CA co-assembly is unclear. In this work, we use Coarse-grained Molecular Dynamics (CGMD) simulations to elucidate the underlying molecular mechanism of capsid formation, including the crucial role played by IP6. We show that IP6, in relatively small quantities at first, promotes curvature generation by trapping pentameric defects in the growing lattice and shifts assembly behavior towards kinetically favored outcomes. Our analysis also suggests that IP6 can stabilize metastable capsid intermediates and can induce structural pleomorphism in mature capsids.

**Teaser:** Computer simulations reveal that IP6 promotes fullerene-like capsid formation by stabilizing high curvature regions of the capsid.

## Introduction

The HIV-1 replication process requires budding of immature viral particles that subsequently undergo a crucial structural rearrangement called maturation.(*1, 2*) HIV-1 maturation is a string of biochemical and morphological changes induced by proteolytic processing of Gag by viral protease (PR),(*1, 3*) leading to self-assembly of released capsid protein (CA) into a cone-shaped capsid(*4*) (mature core) encasing a dense RNA/nucleocapsid (NC) complex.(*5*) The mature capsid executes several critical functions during reverse transcription (RT)(*6*), cellular trafficking(*7*) and nuclear import.(*8, 9*) Thus, mature capsid generation is essential for HIV-1 infection, with failure resulting in loss of viral infectivity.(*10–12*)

The CA monomer contains two independent domains, the N-terminal domain (NTD) comprising seven α-helices and a β-hairpin and the C-terminal domain (CTD) comprising four α-helices, that are connected by a flexible linker.(*13*) Interestingly, CA can adopt a dimeric form (Fig. 1a) in solution due to interactions between CTD helices(*13, 14*) and exists in a dynamic monomer/dimer equilibrium.(*14*) CA can self-assemble into a wide variety of morphologies while a typical capsid is a closed fullerene shell (Fig. 1d), composed of hexamer and pentamer building blocks (Fig. 1c) with more than 1000 copies of CA.(*4, 15*) Typical HIV-1 virions contain 2500-5000 Gag monomers in a non-trivial and highly crowded environment.(*16–19*) Therefore, a mature capsid, which contains only half of the CA present in an immature virion, exists in equilibrium with solution state CA. As the assembly condition is sensitive to variation in CA concentration, pH and salt concentration, biochemical reconstitution of CA assembly has been particularly challenging.(*20*) Under favorable conditions, a mixture of cone and cylinder formation was observed in most of the previous *in vitro* studies.(*21, 22*) Cryo-EM and image reconstruction studies have identified several key structural motifs for capsid assembly that involve interactions between the NTD/NTD, NTD/CTD and CTD/CTD interfaces of CA.(*23, 24*) Characterization of CA tubular assemblies showed that the NTDs form a hexagonal arrangement while inter-hexamer CTD interfaces connect adjacent assembly units.(*23*) Based on various *in vitro*(*23, 24*) and intact virion(*25*) analyses, the atomic structure of mature cores, following a fullerene cone model, with exactly 12 pentamers(*22, 24, 26*) in agreement with Euler’s theorem, has also been determined. Recent X-ray crystallography studies reveal that a narrow pH gated pore with a ring of R18 at its center is distributed throughout the capsid surface.(*27, 28*) This ring of R18 is essential for RT and avidly binds several host cell metabolites.(*27*) Interestingly, HIV-1 packages more than 300 ions of negatively charged IP6 per virion,(*29*) which has been found to influence immature Gag assembly.(*29, 30*) Pioneering work by Dick *et al.* demonstrated that IP6 is also critical for enhanced assembly of cone-shaped mature cores.(*31*) IP6 has been shown to interact with the ring of R18 and removal of electropositive repulsion in the central pore by R18A mutation precludes viral infectivity.(*31*) Therefore, it is evident that IP6 plays a central role in infectious capsid formation. In a previous work from this group,(*32*) the structural basis for interaction between IP6 and capsid building blocks was illustrated using all-atom (AA) molecular dynamics (MD). Importantly, free energy calculations indicated that IP6 preferentially stabilizes CA pentamers over hexamers. Therefore, it was hypothesized that IP6 promotes capsid formation by stabilizing pentamers, which are necessary for inducing curvature in the capsid. However, the detailed molecular mechanism of IP6 and CA co-assembly and several important questions regarding the kinetic mechanism of pentamer inclusion and curvature generation during the HIV-1 self-assembly remain elusive.

**Figure 1.**
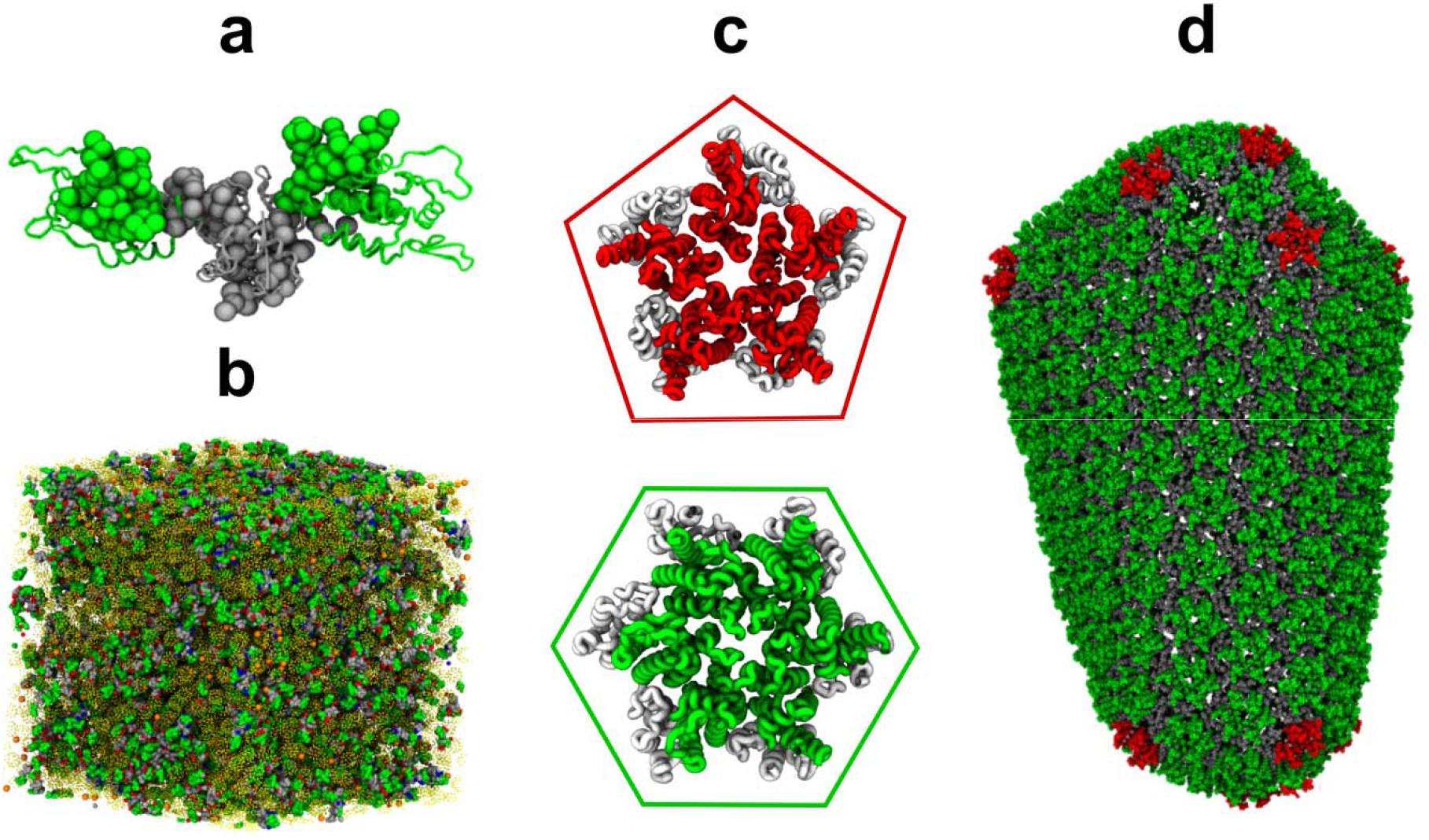
CG model of HIV-1 CA dimer and CGMD simulation setup. (a) Overlay of a CG CA dimer (NTD green and CTD gray) over an all-atom CA dimer (PDB 2ML8).(*14*) Shown in this figure are the CG sites or “beads”. (b) Snapshot of an initial simulation setup; CA dimer (green, gray beads and blue/red for active/inactive), IP6 (orange beads), inert crowder (transparent yellow). (c) Quasi-equivalent pentamers (PDB 3P05) and hexamers (PDB 5HGL) after ref 15 and 26 respectively. CTDs are gray and NTDs are red for capsid pentamer and green for hexamer. (d) Mature core as made of pentamer (red) and hexamer (green) building blocks.

All-atom simulations offer complementary insight to experimental studies. AAMD simulations of full-capsid have revealed several key physical and chemical properties of capsid dynamics and structure,(*33, 34*) nonetheless, these techniques are computationally demanding and, therefore inadequate for studying the large-scale functional dynamics of capsid assembly processes that occur over long time scales. Alternatively, coarse-grained (CG) approaches reduce system complexity to a simpler molecular description while retaining the ability to predict emergent behavior.(*35*) While caution should always be exercised when considering large-scale molecular assembly, general agreement with experimental results is often considered a reliable indicator of CG model fidelity. Previous CG simulations(*36, 37*) have offered valuable insight into the effects of CA conformational freedom, concentration, and molecular crowding on self-assembly, thus explaining the initial stages of capsid growth. In this work, however, we extend our prior CG model(*36*) to include, and investigate, the effects of IP6 on HIV-1 capsid formation. In particular, we examine the kinetic pathways promoted by IP6 that lead to capsid morphogenesis. Importantly, we observe that IP6 is essential for the formation of high curvature lattice regions representing kinetically favored intermediates. We therefore propose that a physiological role for IP6 is to induce lattice curvature by stabilizing a small population of CA pentamers during the early stages of the capsid self-assembly. Subsequent to that process, a larger number of IP6 ions then intercalate into the formed or nearly formed capsid lattice to add further stabilization.

## Results

### CA assembly behavior in the absence of IP6

HIV-1 CA monomers consist of two globular domains (NTD and CTD) connected by a flexible linker. The monomer is conformationally flexible and adopts an assembly competent (i.e. mature-like) state ([CA]_+_) with only 5-10% probability in solution.(*14*) To emulate this behavior in our CG simulations, CA was partitioned into assembly competent ([CA]_+_) and assembly incompetent ([CA]_-_) populations and all free CA were periodically assigned to either the [CA]_+_ or [CA]_-_ state with a fixed [CA]_+_ probability; the state switching rate was chosen to maintain assembly relevant correlation times (conformational switching interval), i.e., inter-domain motions. This approach is the so-called “ultra” CG (UCG) model in which various physical or chemical processes can implicitly be overlaid on the kinematic motion of the CG particles in order to describe (and include) features that are impossible to include at the explicit CG resolution of the model. UCG modeling can be exceptionally powerful and, in this case, has enabled the simulation of HIV-1 capsid formation from more than 1000 proteins. As noted earlier, herein the UCG model for HIV-1 capsid has been modified to include explicit CG IP6 anions in the simulation.

Several *in vitro* experiments have demonstrated that, in the absence of IP6, CA predominantly forms hollow tubes made of hexamers.(*24, 31*) While our prior CG model(*36*) of CA predicted cone formation, we tested whether our current modified CG model was also capable of producing tubes, resulting in a mixed population of tubes and cones. To simulate CA assembly, 4 independent trajectories with 600 CA dimers were propagated for 5 × 10^8^ CG steps and 200 mg/mL inert crowder density with [CA]_+_/[CA]_free_ = 15% and conformational switching interval of 5 × 10^5^ time steps. The CA concentrations in our simulations were about 50% of a typical virion due to computational expense. However, by increasing [CA]_+_ to 15% in comparison to our prior studies(*36, 38*) that used [CA]_+_ = 10%, we compensated for the possible undersupply of assembly competent CA. From our CG simulations, we observed 3 tubular structures and 1 truncated cone (Figs. 2a and S1). CA dimers quickly produced metastable trimer of dimer populations (< 5 × 10^6^ CGMD steps), but nucleation of these trimers was slow and did not occur before 10^7^ CG steps. The growing lattice curled into a hollow cylinder and grew rapidly by CA dimer association at both ends generating a sheet of hexamers (Fig. 2a). After 3 × 10^8^ CG steps, pentamer defect incorporation at the edge of the lattice was observed forming a narrow end (Fig. S1). Interestingly, CA self-assembly at the exposed lattice edges was dynamic and exhibited a degree of local remodeling by “error correction”. The other end of the CA tube continued to grow until the population of assembly competent CA was exhausted in the simulation box. Snapshots taken at various stages of CA assembly in the absence of IP6 are shown in Fig. 2a and the full process is shown in the movie S1. As shown in Fig. 2b, the final CA tube containing 845±40 CA monomers was primarily composed of hexamers (126 ± 7). The diameter of the final structure was 38 ± 6 nm, which is consistent with tube diameters from *in vitro* experiments.(*23*) The rarer appearance of truncated cones (Fig. S1) reflects the ability of our CA model to assemble spontaneously into both helical tubes and cones with a preference for tubular structures in the absence of IP6. These results are reproducible in additional simulations albeit with a statistical spread.

**Figure 2.**
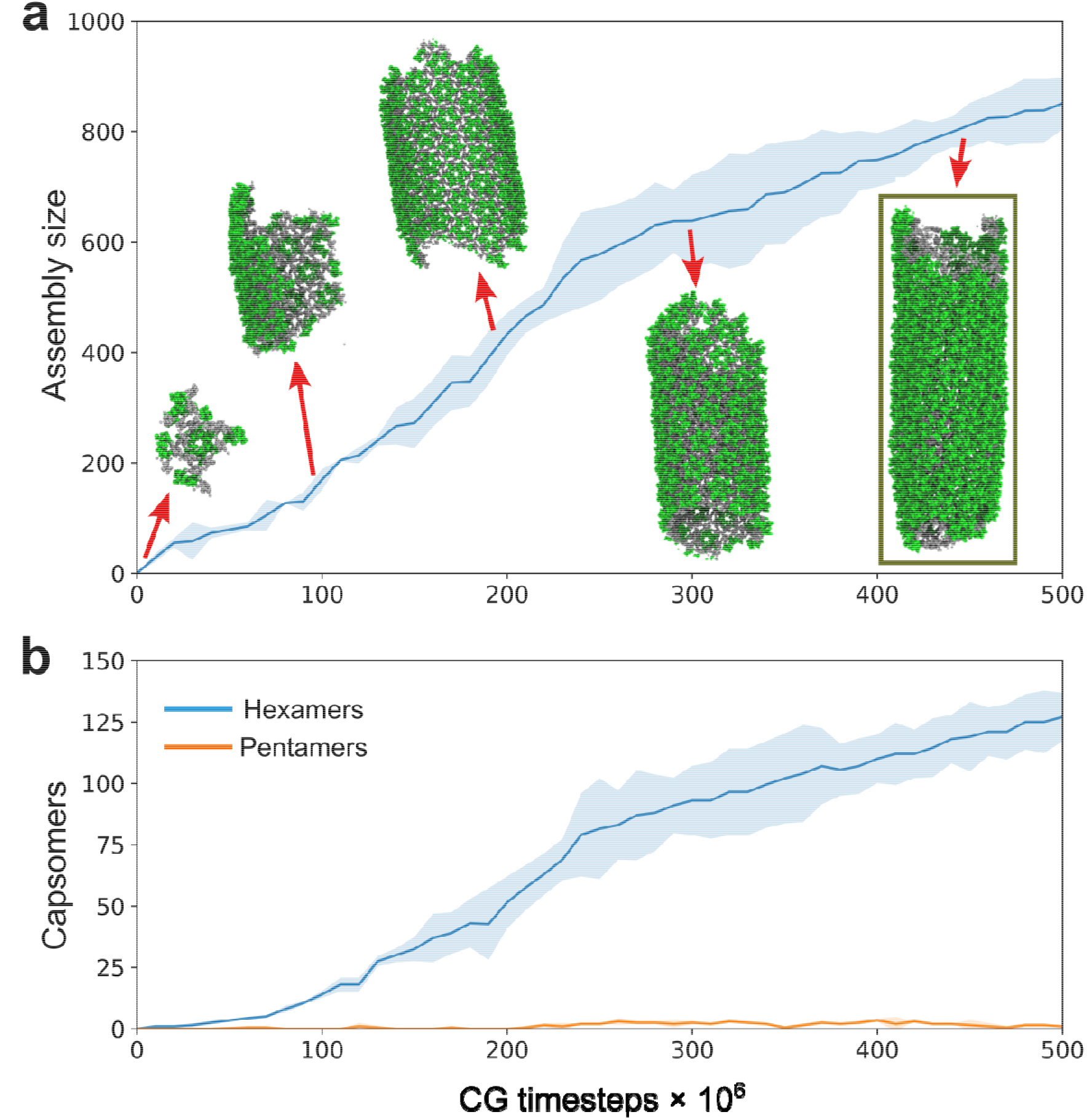
CGMD simulations of CA dimer assembly in the absence of IP6. (a) Assembly time-series plot that depicts cluster size as a function of CG time. CA dimers preferably assemble into tubular structures in the absence of IP6 (CA NTDs are green and CTDs are gray). Simulation snapshots taken at,,,,CGMD time steps. (b) Time-series profile of the total number of hexamers and pentamers as a function of CG time step. In both panels, the shaded region represents the standard deviation across all four replicas.

### Molecular mechanism of IP6 mediated CA assembly

Recent studies have shown that the negatively charged host metabolite IP6 can stabilize the six helix-bundle(*31*) in the immature Gag and is efficiently recruited by HIV-1 virions.(*39*) After proteolysis, IP6 is known to stabilize the mature capsid by binding at the central ring of R18.(*32*) IP6 is also critical for enhanced mature capsid formation. (*31*) Thus, IP6 shifts the population of assembled CA morphologies towards mature capsid cones through as-yet unknown mechanisms. A typical mature capsid contains 1000-1500 CA monomers.(*4, 15*) Therefore, we increased the simulation box domain to 90×90×90 nm^3^ with 880 CA dimers to allow the formation of complete capsids. To examine the effects of IP6 during self-assembly, simulations were performed with 4mM CA and 600 IP6 molecules using the aforementioned assembly conditions, including the concentration of inert crowders and the CA conformational switching rate described earlier. Figure 3 presents different stages of IP6 mediated capsid assembly. A small and curled hexameric lattice formed initially with small curvature since the dihedral angle between adjacent hexamer-hexamer planes was close to 180°. This explains the formation of the relatively flatter middle region in the initial stages of capsid assembly (~ CGMD steps). Subsequently, stable pentamers formed at the broad end of the capsid and were stabilized by IP6, imparting curvature to the capsid shell. The growth of the capsid lattice followed a sigmoidal profile, a characteristic also observed experimentally.(*40*)

**Figure 3.**
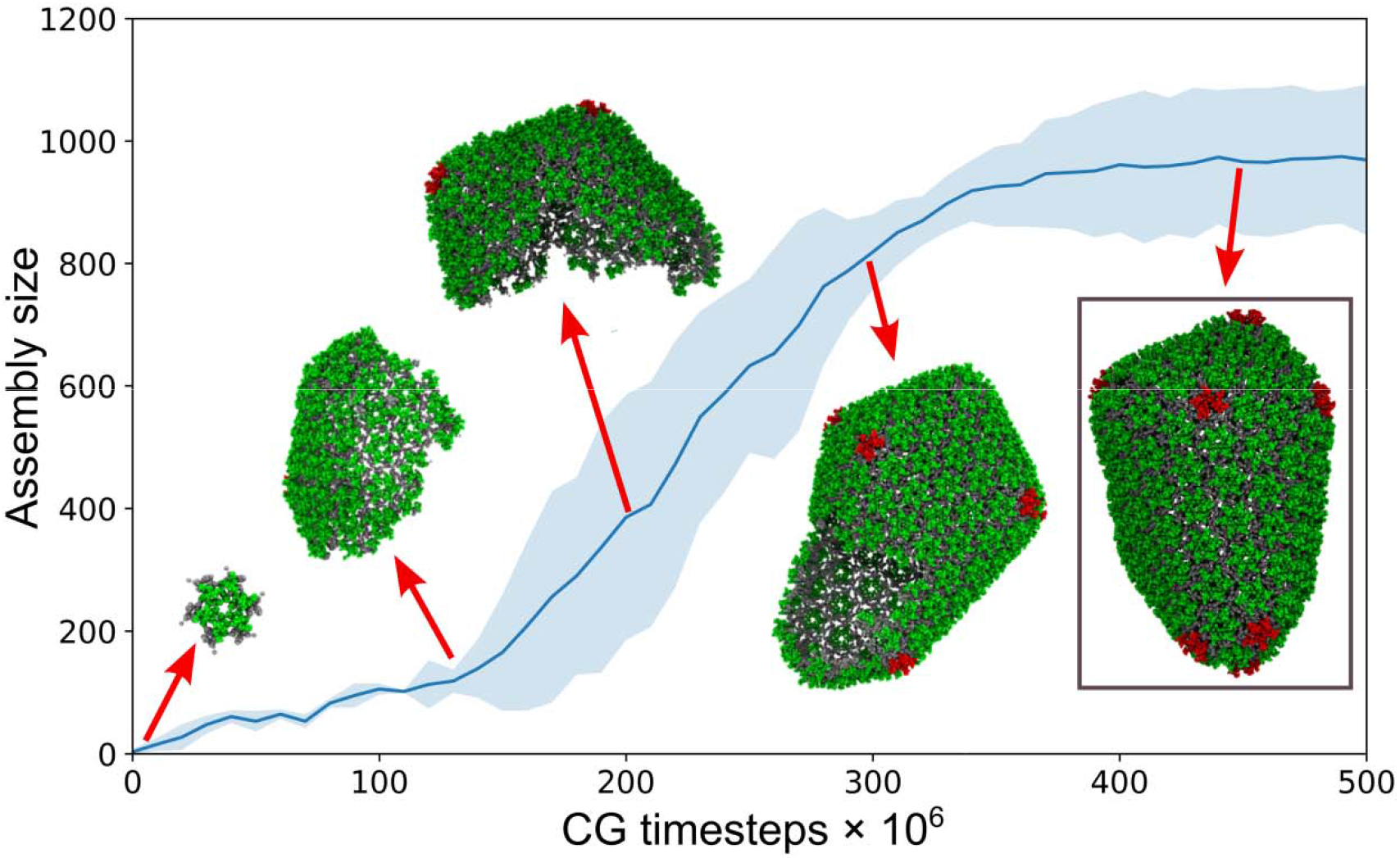
CGMD simulations of CA dimer assembly in the presence of IP6. (a) Time-series plot of assembled lattice size as a function of CG time step. CA dimers assemble into a conical capsid in the presence of IP6 (CA hexamer NTDs are green, pentamer NTDs are red and all CTDs are gray). Simulation snapshots taken at,,,,CGMD time steps. The shaded region represents the standard deviation across six simulation replicas.

Interestingly, the narrow end of the cone formed last in our simulations where five proximal pentamers emerged, leading to the high-curvature cap shown in Figure 3 and movie S2. For comparison, our prior CG model(*36*) in the absence of IP6 tended to form the narrow high curvature cap first. In the present case, the complete capsid contained 978±121 CA subunits and was observed after CGMD steps. We analyzed time series profiles of CG trajectories in Figure 4, in which we find that the complete capsid core is composed of 151±21 CA hexamers and exactly 12 pentamers.

**Figure 4.**
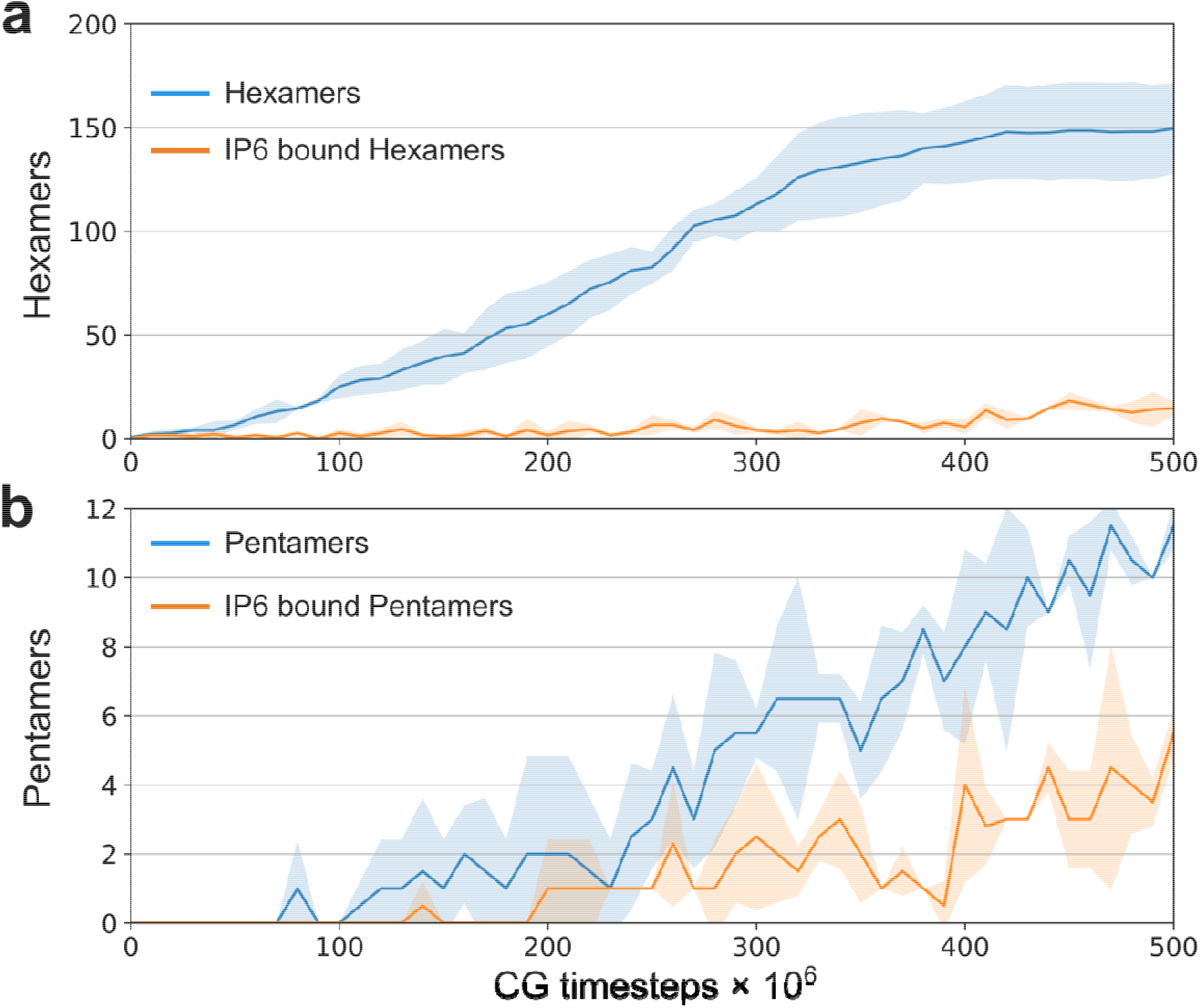
Assembly time-series profiles illustrate the preference of IP6 towards pentamers over hexamers. (a) The total number of hexamers and the total number of IP6 bound hexamers in the capsid as a function of CGMD time, (b) The total number of pentamers and the total number of IP6 bound pentamers in the capsid as a function of CG time. In both panels, the shaded region represents the standard deviation across six simulation replicas.

To further understand the role of IP6 in CA assembly kinetics, we counted the number of IP6 bound capsid hexamers and pentamers as the simulation progressed. While IP6 bound with only 15-20% of the hexamers, by contrast at least 50% of the pentamers were bound with IP6 (Fig. 4a & b), indicating that IP6 shows preference towards pentamers in our simulations even under such complex conditions.

### IP6 accelerates CA assembly rate and stabilizes regions of high curvature

To further investigate the precise behavior of CA assembly, we compared the early stages of assembly in the absence and presence of IP6. We performed 10 additional short simulations, 5 each for CA without IP6 and CA with IP6 starting from a pre-nucleated hexameric cluster with 6 CA dimers. We depict the assembly rate of CA in Figure 5 and analyze the two cases. In the presence of IP6, the assembled clusters contain 150 ± 44 CA monomers after CGMD steps. During the early stages of assembly, IP6 binds with the nucleated cluster and accelerates CA assembly compared to that of the IP6-absent condition. While local remodeling at the growing lattice edges is a key feature of capsid assembly, we observed that IP6 binding stabilizes defects in capsid intermediates, thereby curtailing extensive error correction and facilitating CA association.

**Figure 5.**
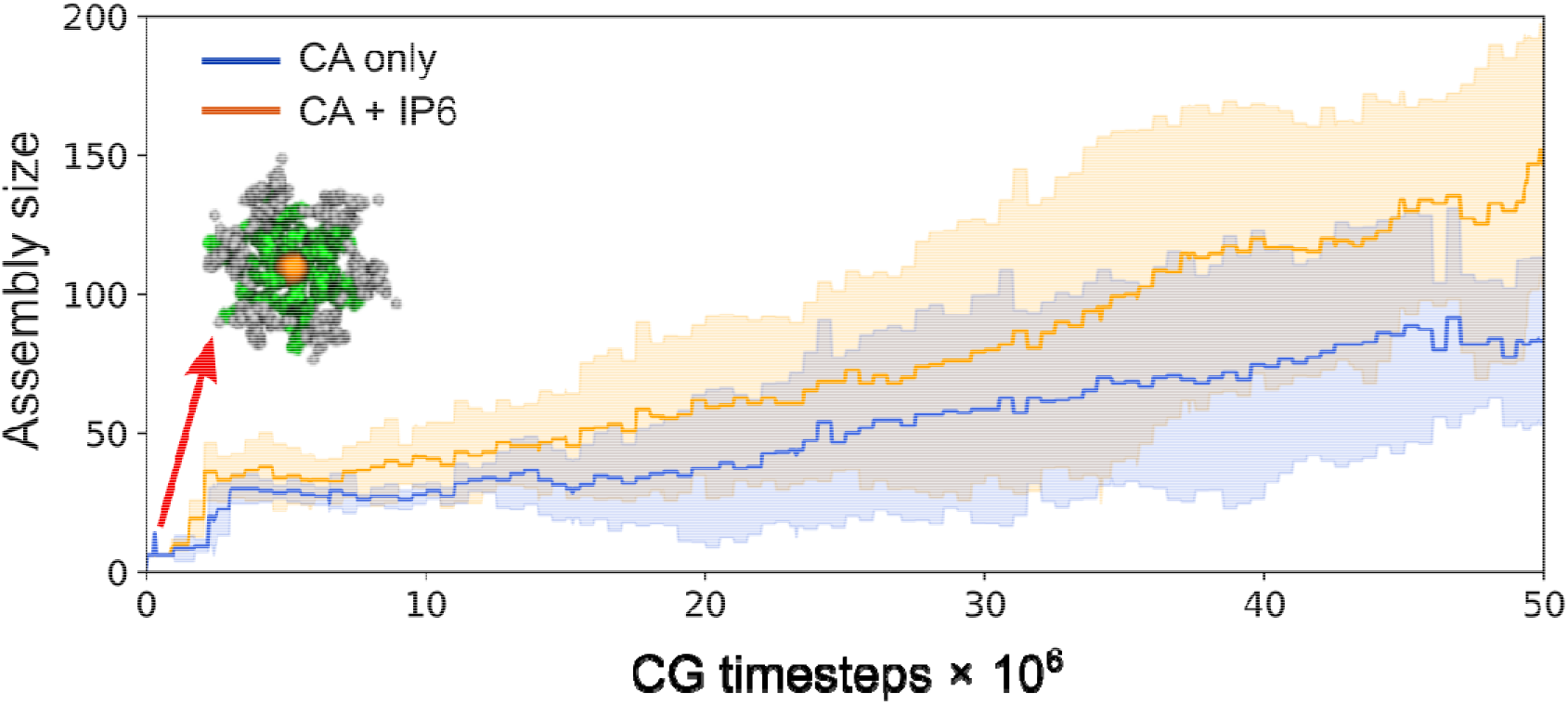
IP6 accelerates CA assembly rate by kinetically trapping capsid intermediates (color scheme is same as Figure 3. IP6 is orange here). Time series profiles of cluster size as a function of CGMD time step in the presence and absence of IP6. In both panels, the shaded region represents the standard deviation across five simulation replicas.

It has been noted that HIV-1 capsids possess a caged polyhedral arrangement composed of CA hexamers and exactly 12 pentamers.(*4, 24*) These pentamers introduce curvature on the capsid and are necessary for capsid closure. On the basis of 2.5 Å X-ray crystallography structures, hexamers and pentamers were determined to be quasi-equivalent(*15*) with the positively charged R18 sidechains throughout the central pore juxtaposed more closely in the pentamer compared to the hexamer, yielding stronger electrostatic repulsion in the former. Pentamers are considered defects and are disfavored compared to regular hexamers. Our CA simulations also show a certain degree of error correction where defective pentamers are annealed into hexamers. Intriguingly, at the end of the CGMD runs when the capsid had formed, the high curvature regions both at the broad and narrow ends of the capsid had more IP6 population than the comparatively flatter middle region as shown in Figure 6.

**Figure 6.**
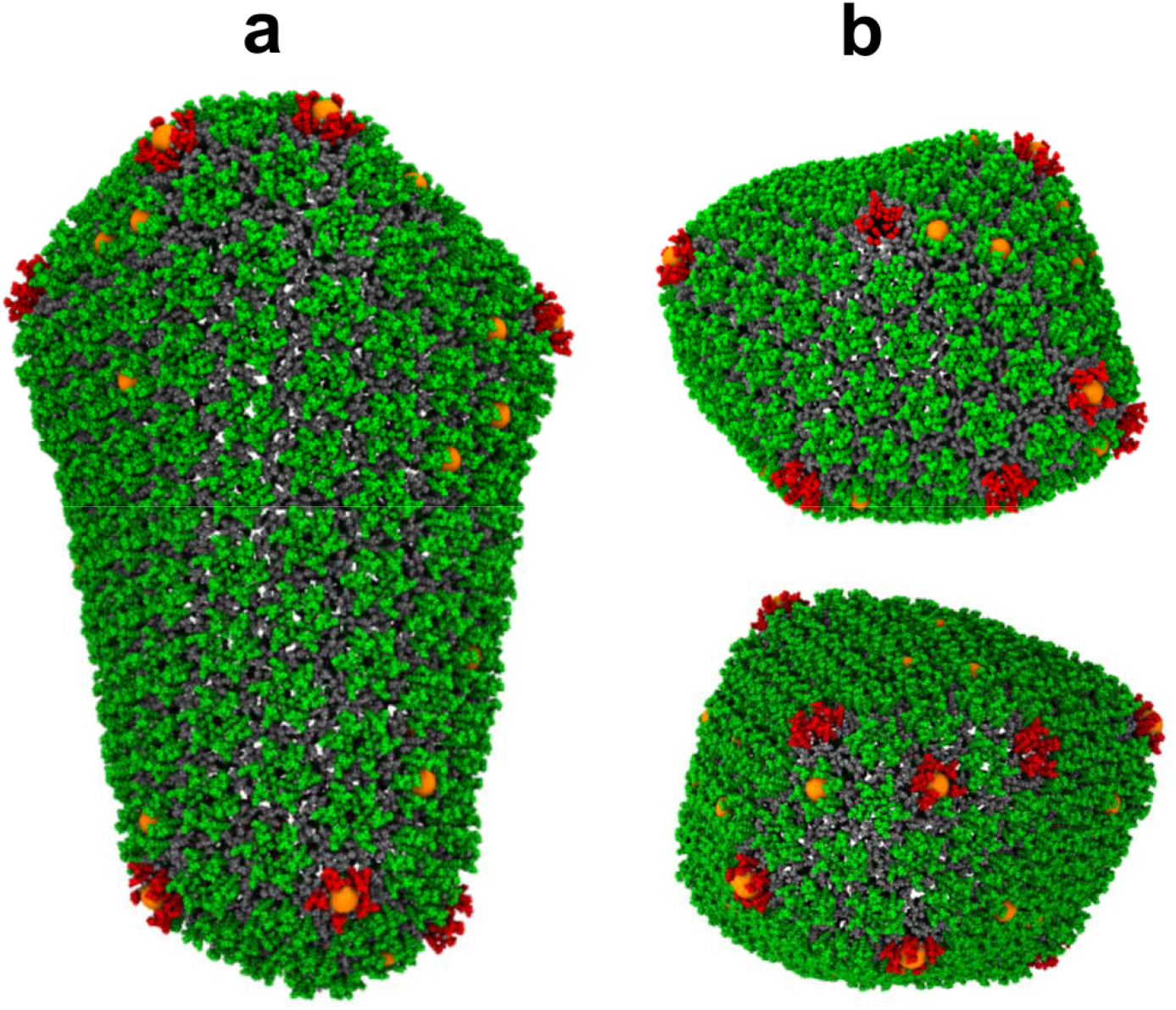
CGMD simulations reveal that IP6 stabilizes the capsid by binding at regions of high curvature. (a) Side view of capsid (color scheme is same as Figure 3). (b) Top and bottom view.

We observed that IP6 prefers binding to the regions of high curvature and are responsible for lattice curvature generation. It should be noted that only 15 to 35 IP6 ions intercalated with the capsid shell during CA assembly in the CGMD runs but IP6 binding continued even after full capsid formation (Fig 4). Recent experimental studies show that the narrow end of the HIV-1 core may not close if the conditions are not favorable.(*41*) However, all capsids generated during our simulations closed completely suggesting that IP6 assists in closure by stabilizing pentamers which accommodate curvature at regions of high stress. This is largely an effect occurring on the kinetic landscape along the capsid formation pathway, while the thermodynamic minimum generally involves the fully formed capsid with significantly more IP6 ions intercalated into its CA lattice.

### IP6 stabilization of capsid intermediates determines mature virion morphology

The fullerene core is a defining feature of HIV-1 maturation.(*3*) However, it is not unusual for HIV-1 viral capsids to display morphological diversity.(*21, 25, 40, 42, 43*) Depending on the spacing between pentameric defects, capsid cores adopt a wide distribution of cone angles thereby producing different morphologies. Analogously, the variable distribution in capsid sizes is suggestive of a robust assembly process. Figure 7 presents different morphological outcomes during our simulations of CA self-assembly with IP6. We observed that curvature generation in capsid intermediates was always dictated by stabilization of pentameric defects by IP6. Unusually early incorporation of these pentameric defects resulted in smaller cores (Figure 3 and 7b), whereas more timely pentamer incorporation into the forming extended hexagonal lattice sheet led to appropriate curvature generation for relatively larger mature core formation as shown in Figure 7a,c,e. Although rarer than the regular conical capsid, a pill-shaped core (Fig. 7d), capped by the introduction of six pentameric defects at both ends, was also observed during our simulations. It is noteworthy that our simulations even reproduce this unusually shaped capsid, as well as other structures, seen in cryo-ET studies of actual mature HIV-1 virions.(*25*)

**Figure 7.**
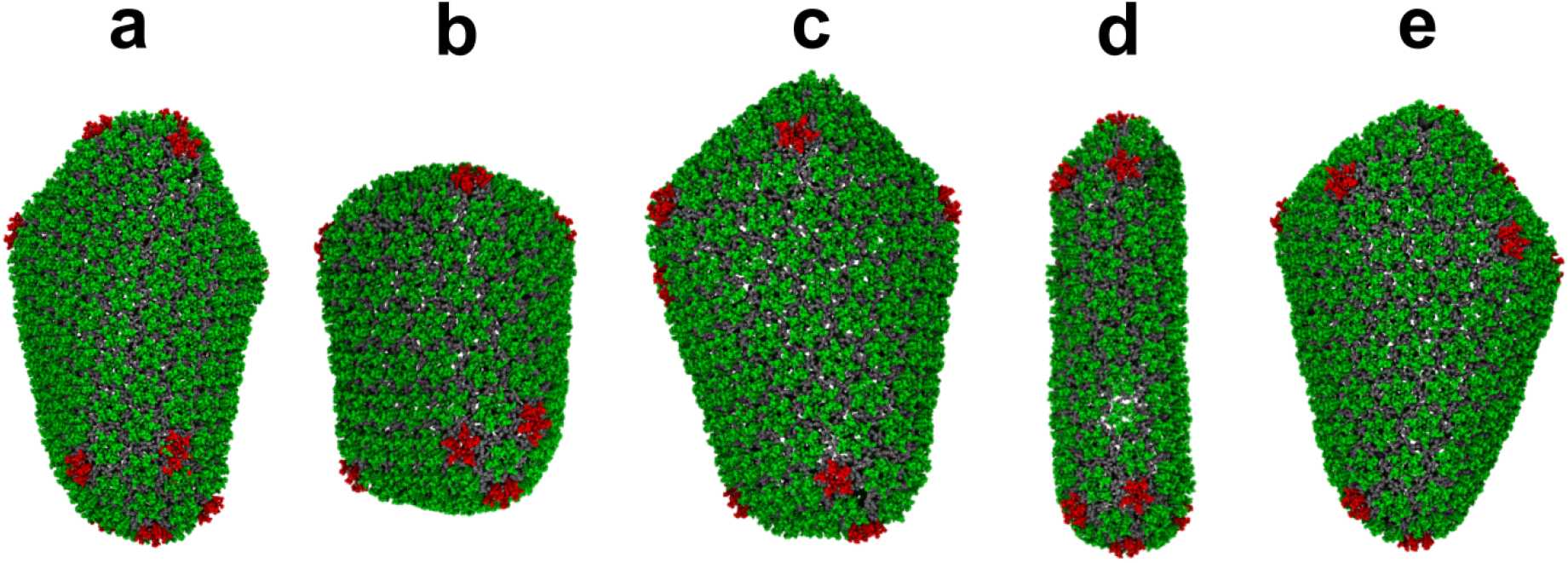
IP6 promotes capsid pleomorphism. (a-e) Snapshots of final structures from five additional CGMD simulations in the presence of IP6 (CA color scheme as in Figure 3). IP6 molecules stabilize lattice curvature inducing the formation of kinetically trapped structural outcomes.

## Discussion

We have used CGMD simulations to elucidate the critical role of IP6 during HIV-1 capsid self-assembly. Our CG simulations capture assembly behavior at time and length scales relevant for viral capsid morphogenesis and they generate virion relevant morphologies, thus demonstrating that CG simulations can provide crucial insight into CA assembly kinetics. While the RNA complex is known to promote capsid nucleation, (*26*) it should be noted that the simulations presented here assume that RNA is not essential for capsid formation, which is in agreement with experiments that indicate cone formation can be achieved in the absence of RNA, although we expect RNA to also accelerate CA assembly.(*44*) In the absence of IP6, we find that mature CA assembly is self-regulated through efficient error-correction, leading to tubule formation. This ensures increased contact among CA monomers and removal of transient pentamer defects by local remodeling, suggesting that CA assembly nominally proceeds through a thermodynamically-driven pathway producing near-equilibrium structures. However, depending on the local [CA]_+_ concentration and crowding, lattice growth can be too rapid for error correction, leading to conical structures with stable pentamers embedded within the lattice as seen occasionally in our simulations and select *in vitro* experiments.(*22*) This explains the primary formation of cylinders and the rarer occurrence of cone-shaped mature cores in the absence of IP6.

The morphological transition from an immature spherical Gag lattice to the mature core within HIV-1 virions is not well understood but there are two competing models for this transformation: the disassembly/reassembly model(*45*) and the displacive mechanism.(*46, 47*) Evidence in support of the disassembly/reassembly model includes several structural studies demonstrating distinct lattice spacings and different protomer-protomer contacts between the immature and mature shells.(*5, 48*) Alternatively, the transition of CA-SP1 mutants with cleavage defects into mature cores supports the displacive model.(*47*) Recent *in vitro* experiments and phenomenological computer simulations indicate a sequential combination of both displacive and disassembly/reassembly pathways during capsid maturation(*49*) where the growth of nucleated CA proceeds through initial formation of the broad capsid followed by capsid closure at the narrow end.(*50, 51*) Our simulations explore the basis of IP6 mediated capsid growth and closure, which is relevant for both the sequential displacive and disassembly/reassembly pathways and CA assembly under *in vitro* conditions. It has been unclear why viral capsids assume conical morphologies with pentameric defects over more stable cylinders. However, our results show that n the absence of IP6 pentamers are unstable and eventually get annealed into hexamers, as evidenced by fluctuation in pentamer population in our simulations (Fig. 2b). By contrast, in the presence of IP6 a combined effect of accelerated assembly and early pentamer stabilization by IP6 allows for the conical structure to form with the incorporation of exactly 12 stable pentamers. However, we note that, over our simulation timescales, not all pentamers bind to IP6, indicating that only a few pentamers stabilized by IP6 are sufficient to facilitate curvature generation with subsequent pentamer formation and hence the closed capsid formation. This result also explains CA tubule formation under low concentrations of IP6 *in vitro*.(*31*) Cone formation may be difficult to accomplish at low concentrations of IP6 due to the stochastic nature of pentamer trapping by IP6 intercalation. It should be noted, however, that our simulations predict additional IP6 intercalation even after the full capsid formation. Our simulations also suggest a delicate balance between curvature generation and lattice growth during CA self-assembly. Therefore, depending on the extent of capsid growth achieved before the inclusion of pentamers necessary for capsid closure, mature capsids may contain 1000-1500 CA monomers, explaining the heterogeneity in capsid shape and size.(*21*)

Interestingly, a single CA protein is capable of wide conformational flexibility, as evident by the range of morphologies with varied curvature in expressed and purified capsids.(*24, 52*) Modeling studies of the HIV-1 capsid revealed that pentameric declination hinges upon flexibility in the CTD-CTD dimer interface,(*15*) whereas the NTD-CTD interface is believed to modulate variable lattice curvature in both hexamers and pentamers by limiting the movement of CA-CTD with respect to CA-NTD.(*42*) Nonetheless, the free energy penalty for sharp lattice curvature formation, as seen in some of non-canonical capsid morphologies, can be prohibitive as evident from the ubiquitous appearance of CA cylinders (tubules) in the absence of IP6.

In our simulations, IP6 ions stabilize capsid intermediates, thereby enabling the formation of high curvature regions and relatedly promoting non-canonical core formation. We thus speculate that small molecules that target IP6 binding sites may preclude HIV-1 viral infectivity by inducing early capsid closure through rapid incorporation of pentamer defects or by over-stabilizing the capsid.(*38*) Considering the hierarchical nature of capsid assembly, our results underscore the importance of the IP6 binding site as a target of opportunity for the development of potential maturation inhibitors and can provide valuable insight into that effort. It is of note that the need for efficient and authentic assembly imparts considerable genetic fragility on HIV-1 CA.(*53*) However, the current CG model is derived from experimental phenomenology which is incapable of providing detailed chemically-specific insights or interactions and, therefore, is unable to explore CA sequence mutation related variation in assembly properties. Further simulations that probe the basis of assembly behavior of CA variants will likely require incorporation of all-atom statistics recapitulating the configuration space of protein mutation. Additionally, the CG interactions between the ring of R18 and IP6 can be further modified using, e.g., coarse-graining technique such as relative entropy minimization(*54*) to approximate the configurational dependence of the potential of mean force by parameters that accurately reproduce structural features at all-atom resolution. The recruitment of nucleotides into the positively charged cavity of R18 is also essential for reverse transcription. Since IP6 plays an important role in stabilizing the ring of R18, additional CG simulations that target the role of IP6 in nucleotide recruitment and import at the virion level will be valuable in an effort to understand the conformity of the capsid pore as a potential drug target.

## Materials & Methods

### CG model details

The NTD and CTD of a CA CG monomer were represented as independent elastic network models (ENMs) containing Cα atoms from CA α–helices, while a weak ENM maintained inter-domain connections. The CG NTD was generated from PDB 3H4E(*55*) using average NTD Cα positions as the location of CG particles. The potential energy of the bonds (*U_bond_*) was given by:

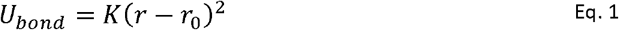

where *r* is the separation distance, and *r*_0_ is the computed average distance (full data in Table S2).

To preserve the local helix structure, an elastic network connected CG particles within an α–helix with harmonic spring constant *K* = 10 kcal/mol/Å^2^ while the relative positions of α–helices were preserved by connecting each CG particle with its nearest neighbor from every other helix with harmonic spring constant *K* = 10 kcal/mol/Å^2^. The complete CA CG ENM network is summarized in Table S1. CA CTD was constructed from chain A of PDB 2KOD(*56*) using the same method and the two CTDs in the CA dimer were connected with an ENM (*K* = 10 kcal/mol/Å^2^). The CA protein assumes an assembly competent (or active) and assembly incompetent (or inactive) state depending on its conformational flexibility controlled by the flexible linker region. Therefore, an additional weak ENM (*K* = 0.01 kcal/mol/Å^2^) was employed to connect CA NTD and CTD. It should be noted that the intent of the model is to maintain the internal shape of the CA dimer to study its self-assembly while reproduction of its detailed structure is not intended here. We used an inert molecular crowding agent to represent the mass and excluded volume of Ficoll70.(*40*) Ficoll70 was modeled using two independent spheres of CG sites connected by ENMs (*K* = 10 kcal/mol/Å^2^), each consisting of 42 CG beads and ≈ 5.1 nm diameter. Ficoll70 CG beads had a default excluded volume separation of 1 nm, an effective radii of 0.5 nm. To capture the molecular shape of CA, CG monomers were overlapped with PDB structures 3H4E(*55*) and 3P0A(*15*) with excluded volume separations adjusted to maintain the relative separation between CG particles. A soft cosine excluded volume potential with an effective radius was used to prevent unphysical overlap between CG beads:

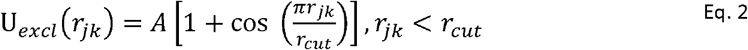

Here, *r_jk_* is the distance between CG particle centers with *r_cut_* = 20Å the onset of excluded volume repulsion for specific CG sites *j* and *k.* The value of *A* is 10 kcal mol^-1^Å^-1^ in all cases.

The CG model also incorporated well-conserved and important CA protein/protein interfaces that were previously determined by centroid analysis of experimental structures 3H4E and 3P0A.(*15, 55*) “Ghost” CG beads were embedded into the CA CG structure at these selected centroid locations (residue 51 and 57 to the NTD, residue 63 and 204 to the CTD)) to function as binding regions in assembly and interacted only with non-ghost CG particles of adjacent monomers. The interaction between the “ghost/non-ghost” particles were represented by attractive interactions which can be enabled and disabled, generating assembly competent ([CA]_+_) and assembly incompetent ([CA]_-_) CA populations. The binding potential was given by a double-Gaussian:

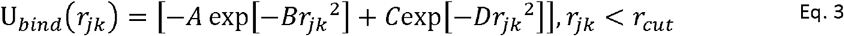

Where *η_k_* is the separation distance between CG particles *j* and *k* with *r_cut_* = 30 Å. The parameters *A* = 1.1 kcal mol^-1^, *B* = 0.1 Å^-2^, *C* = 2.0 kcal mol^-1^, and *D* = 0.01 Å^-2^ describe the depth and width of the long-range and short-range Gaussian interactions, respectively. Full details of the CG models for CA and the inert crowding agent are described in ref 36. Aditionally, we used virion relevant assembly conditions and an explicit IP6 to produce mature-style capsids over the simulation length examined. We generated a single site CG model for IP6 using a linear center-of-mass mapping. Interactions between CA and IP6 were represented by an attractive interaction, with the locations of energy minima parameterized using the CG model of the full capsid. The CA self-assembly process is sensitive to the specifics of the local environment. In addition to that, accurate representation of the weak binding effects between proteins and the dynamics of IP6 binding with the growing lattice in virion-relevant conditions renders CG parameterization challenging. Therefore, consistent with the CG CA model, we modeled the attractive interaction between the R18 residue of CA and IP6 using a Gaussian interaction.

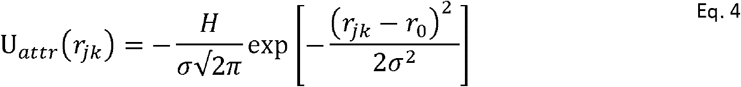

Here *r*_0_ = 8.3 Å is the location of the energy minima determined from the average distance between the vertex and the centroid of the R18 ring within hexamers and pentamers of a preassembled CG capsid (Fig. S2). The parameter *H* = 2.8 kcal Å mol^-1^ and *σ* = 0.4 Å describe the depth and width of the Gaussian interaction, respectively, and were determined by extensive CGMD simulations of IP6 binding to a pre-assembled capsid (Fig. S2). For simplicity, we used parameters to identify the minimum required CG binding energy for at least 50% IP6 occupancy within R18 rings throughout the pre-assembled capsid after 2 × 10^8^ CGMD timesteps.. The mass of each CG particle in CA, inert crowder and IP6 were set to 10 Da, 22.6 Da and 33 Da (≈ 20 fold reduction) respectively to access longer time-scales in the CGMD simulations. All simulations were prepared using Moltemplate.(*57*)

### CGMD simulations

For the CA self-assembly simulations, 1232 monomers, 1700 inert crowders were randomly distributed throughout a 80×80×80 nm^3^ cubic box with periodic boundaries; four replicas were simulated. Another distinct system using a 90×90×90 nm^3^ cubic box with periodic boundaries and with 1760 CA and 2496 inert crowders was prepared for comparing the assembly rates of CA; five replicas were simulated. For the CA and IP6 coassembly simulations, a random distribution of 1760 CA, 600 IP6 molecules and 2496 inert crowders were prepared in a 90×90×90 nm^3^ cubic box with periodic boundaries; six replicas were simulated. All CGMD simulations were performed using LAMMPS 21Jul2020.(*58*) All simulations were performed in a constant NVT ensemble using a Langevin thermostat(*59*) at 300 K with a damping period of 100 ps. The correlation times of solution-state CA inter-domain motion is 2-5 ns. With CG timestep τ = 10 fs, 5 × 10^5^ τ corresponds to 5 ns CG time. Therefore, [CA]_+_/[CA]_-_ the UCG state switching was attempted every 5 × 10^5^ τ For all simulations, a swarm of 50 parallel independent trajectories were propagated for 10^7^ τ initially to expedite the identification of a nucleated cluster. The trajectory yielding a nucleated cluster (size < 12 CA monomers) was used to initialize the production runs. Simulation snapshots were saved every 10^7^ τ.

### Data analysis and visualization

Graph analysis of assembled lattices was performed using the Python package NetworkX 2.1 (http://networkx.github.io/) using two proximity criteria. Inter-hexameric contacts were defined by the CTD dimer interface, i.e., when the distance between CG types 102/102 were less than 2.5 nm. Intra-hexameric contacts were defined by the distance between CA R18, i.e., when CG types 2/2 were less than 1.4 nm. Time series profiles of extracted clusters were generated using Python package Matplotlib.(*60*) Visualization of extracted clusters were generated using VMD 1.9.3.(*61*)

## Acknowledgements

The authors acknowledge the Texas Advanced Computing Center (TACC) at The University of Texas at Austin for providing HPC resources that have contributed to the research results reported within this paper.(*62*) This research also used the Extreme Science and Engineering Discovery Environment (XSEDE), which is supported by the National Science Foundation grant number ACI-1548562.(*63*)

## Author contributions

M.G., A.J.P and G.A.V. designed research. M.G. and A.J.P performed research. M.G. and A.J.P contributed methods, simulation code, and analytic tools. M.G., A.J.P., and G.A.V. analyzed data. M.G., A.J.P., and G.A.V. wrote the paper.

## Data and materials availability

All data needed to evaluate the conclusions in the paper are present in the paper and/or the Supplementary Materials. All relevant data are available from the authors on request.

