## Supplementary material for "Critical Mechanistic Features of HIV-1 Viral Capsid Assembly": SI-ca-ip6

This PDF includes:

Figs. S1 and S2
Tables S1 and S2

Other Supplementary Materials for this manuscript include the following: Movies S1 and S2 show the CGMD simulation of the process of CA assembly without and with IP6, respectively.

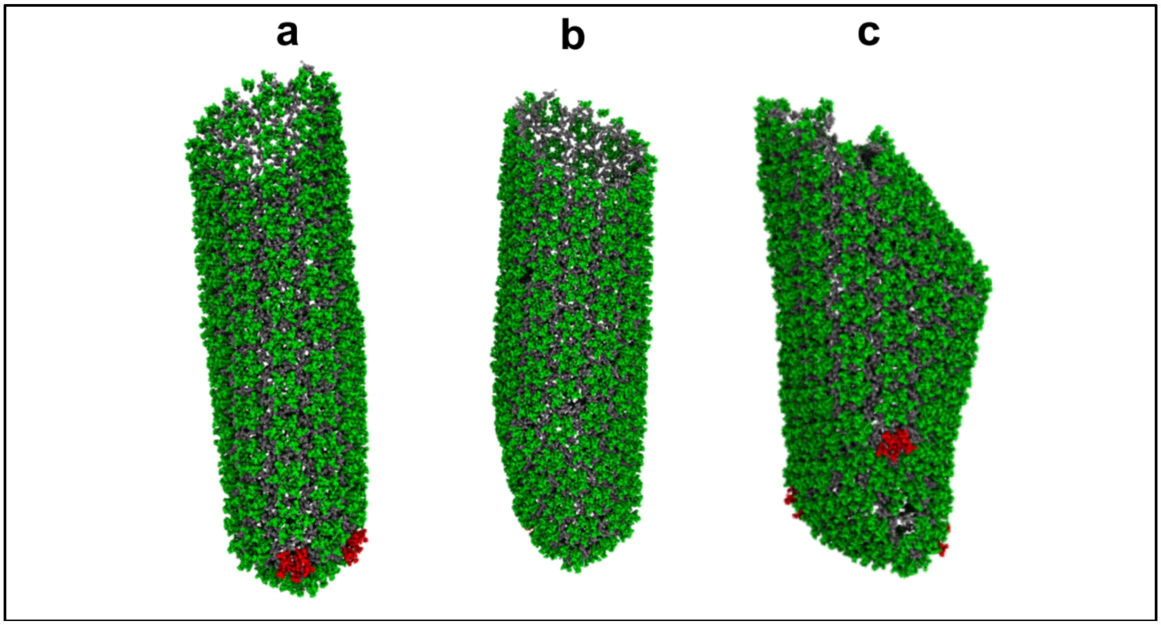


**Fig. S1. CGMD simulations of CA dimer assembly in the absence of IP6.** Snapshots of final structures from three additional CGMD simulations of CA assembly in the absence of IP6 (CA color scheme as in Fig. 3 of the main text). Stable pentamers are embedded in the lattice. In panel (c) a partially conical capsid is formed.

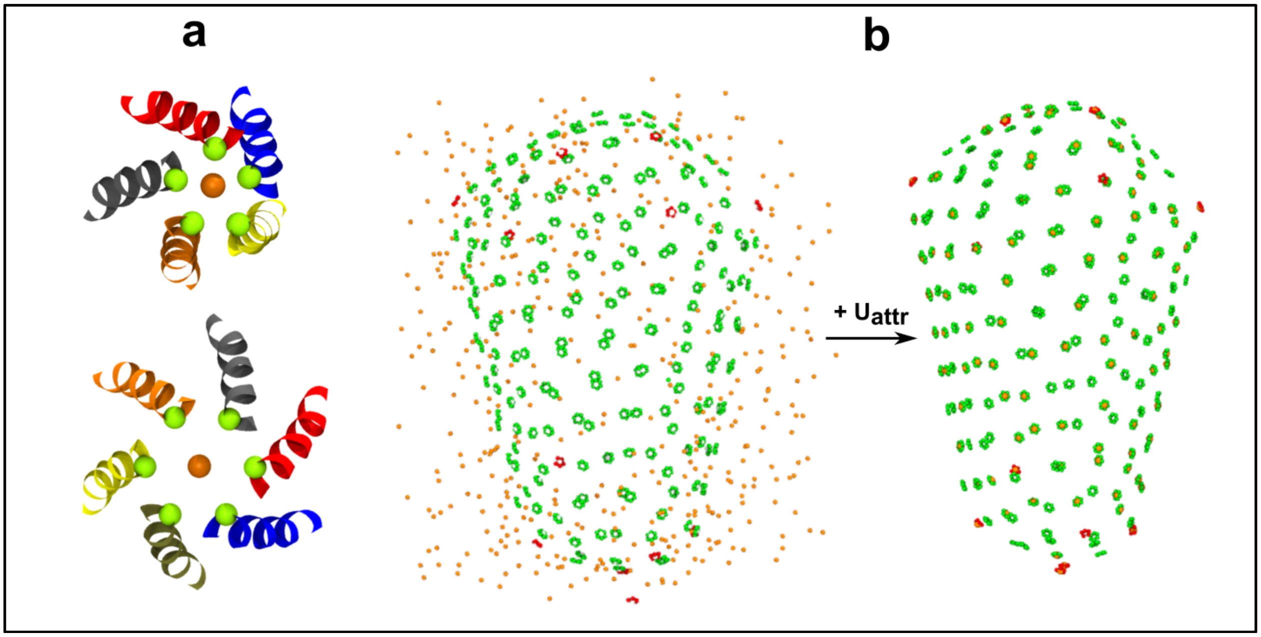


**Fig. S2. Schematic of IP6 binding with pre-assembled capsid, during IP6 parameterization.** (a) CG representation of the R18 ring (green beads) and IP6 (orange bead) in hexamers and pentamers. Helix 1 of the CA hexamer (from PDB 3H4E) and pentamer (from PDB 3P04) shown in distinct colors to highlight the binding location of IP6 in the CG model. (b) IP6 binding to R18 rings throughout the pre-assembled capsid. Only R18 bead (green and red for the hexamer and pentamer, respectively) bound IP6 (orange beads) are shown for clarity.

**Table S1: CA ENM data**

| No. | CG bond | bead 1 | bead 2 |
|---|---|---|---|
| 1 | bond:1CA17CA18 | $atom:1 | $atom:2 |
| 2 | bond:2CA17CA19 | $atom:1 | $atom:3 |
| 3 | bond:3CA17CA20 | $atom:1 | $atom:4 |
| 4 | bond:4CA17CA21 | $atom:1 | $atom:5 |
| 5 | bond:412CA17CA43 | $atom:1 | $atom:22 |
| 6 | bond:426CA17CA55 | $atom:1 | $atom:29 |
| 7 | bond:440CA17CA66 | $atom:1 | $atom:36 |
| 8 | bond:454CA17CA104 | $atom:1 | $atom:57 |
| 9 | bond:462CA17CA111 | $atom:1 | $atom:58 |
| 10 | bond:470CA17CA127 | $atom:1 | $atom:68 |
| 11 | bond:5CA18CA19 | $atom:2 | $atom:3 |
| 12 | bond:6CA18CA20 | $atom:2 | $atom:4 |
| 13 | bond:7CA18CA21 | $atom:2 | $atom:5 |
| 14 | bond:8CA18CA22 | $atom:2 | $atom:6 |
| 15 | bond:413CA18CA43 | $atom:2 | $atom:22 |
| 16 | bond:427CA18CA55 | $atom:2 | $atom:29 |
| 17 | bond:441CA18CA66 | $atom:2 | $atom:36 |

18 bond:463CA18CA111 $atom:2 $atom:58
19 bond:471CA18CA131 $atom:2 $atom:72
20 bond:9CA19CA20 $atom:3 $atom:4
21 bond:10CA19CA21 $atom:3 $atom:5
22 bond:11CA19CA22 $atom:3 $atom:6
23 bond:12CA19CA23 $atom:3 $atom:7
24 bond:414CA19CA43 $atom:3 $atom:22
25 bond:428CA19CA55 $atom:3 $atom:29
26 bond:442CA19CA66 $atom:3 $atom:36
27 bond:455CA19CA104 $atom:3 $atom:57
28 bond:464CA19CA111 $atom:3 $atom:58
29 bond:472CA19CA131 $atom:3 $atom:72
30 bond:770CA19GC51 $atom:3 $atom:131
31 bond:13CA20CA21 $atom:4 $atom:5
32 bond:14CA20CA22 $atom:4 $atom:6
33 bond:15CA20CA23 $atom:4 $atom:7
34 bond:16CA20CA24 $atom:4 $atom:8
35 bond:415CA20CA43 $atom:4 $atom:22
36 bond:429CA20CA55 $atom:4 $atom:29
37 bond:443CA20CA66 $atom:4 $atom:36
38 bond:456CA20CA104 $atom:4 $atom:57
39 bond:465CA20CA111 $atom:4 $atom:58
40 bond:473CA20CA131 $atom:4 $atom:72
41 bond:17CA21CA22 $atom:5 $atom:6
42 bond:18CA21CA23 $atom:5 $atom:7
43 bond:19CA21CA24 $atom:5 $atom:8
44 bond:20CA21CA25 $atom:5 $atom:9
45 bond:416CA21CA43 $atom:5 $atom:22
46 bond:430CA21CA58 $atom:5 $atom:32
47 bond:444CA21CA66 $atom:5 $atom:36
48 bond:457CA21CA104 $atom:5 $atom:57
49 bond:466CA21CA111 $atom:5 $atom:58
50 bond:474CA21CA131 $atom:5 $atom:72
51 bond:21CA22CA23 $atom:6 $atom:7
52 bond:22CA22CA24 $atom:6 $atom:8
53 bond:23CA22CA25 $atom:6 $atom:9
54 bond:24CA22CA26 $atom:6 $atom:10
55 bond:417CA22CA43 $atom:6 $atom:22
56 bond:431CA22CA55 $atom:6 $atom:29
57 bond:445CA22CA66 $atom:6 $atom:36
58 bond:458CA22CA104 $atom:6 $atom:57
59 bond:467CA22CA111 $atom:6 $atom:58
60 bond:475CA22CA138 $atom:6 $atom:79
61 bond:25CA23CA24 $atom:7 $atom:8
62 bond:26CA23CA25 $atom:7 $atom:9
63 bond:27CA23CA26 $atom:7 $atom:10
64 bond:28CA23CA27 $atom:7 $atom:11
65 bond:418CA23CA40 $atom:7 $atom:19
66 bond:432CA23CA55 $atom:7 $atom:29
67 bond:446CA23CA66 $atom:7 $atom:36
68 bond:459CA23CA104 $atom:7 $atom:57

69 bond:468CA23CA111 $atom:7 $atom:58
70 bond:476CA23CA138 $atom:7 $atom:79
71 bond:29CA24CA25 $atom:8 $atom:9
72 bond:30CA24CA26 $atom:8 $atom:10
73 bond:31CA24CA27 $atom:8 $atom:11
74 bond:32CA24CA28 $atom:8 $atom:12
75 bond:419CA24CA40 $atom:8 $atom:19
76 bond:433CA24CA58 $atom:8 $atom:32
77 bond:447CA24CA66 $atom:8 $atom:36
78 bond:460CA24CA104 $atom:8 $atom:57
79 bond:469CA24CA111 $atom:8 $atom:58
80 bond:477CA24CA138 $atom:8 $atom:79
81 bond:33CA25CA26 $atom:9 $atom:10
82 bond:34CA25CA27 $atom:9 $atom:11
83 bond:35CA25CA28 $atom:9 $atom:12
84 bond:36CA25CA29 $atom:9 $atom:13
85 bond:420CA25CA36 $atom:9 $atom:15
86 bond:434CA25CA58 $atom:9 $atom:32
87 bond:448CA25CA63 $atom:9 $atom:33
88 bond:478CA25CA138 $atom:9 $atom:79
89 bond:37CA26CA27 $atom:10 $atom:11
90 bond:38CA26CA28 $atom:10 $atom:12
91 bond:39CA26CA29 $atom:10 $atom:13
92 bond:40CA26CA30 $atom:10 $atom:14
93 bond:421CA26CA36 $atom:10 $atom:15
94 bond:435CA26CA55 $atom:10 $atom:29
95 bond:449CA26CA66 $atom:10 $atom:36
96 bond:479CA26CA138 $atom:10 $atom:79
97 bond:41CA27CA28 $atom:11 $atom:12
98 bond:42CA27CA29 $atom:11 $atom:13
99 bond:43CA27CA30 $atom:11 $atom:14
100 bond:422CA27CA36 $atom:11 $atom:15
101 bond:436CA27CA56 $atom:11 $atom:30
102 bond:450CA27CA65 $atom:11 $atom:35
103 bond:461CA27CA104 $atom:11 $atom:57
104 bond:480CA27CA138 $atom:11 $atom:79
105 bond:44CA28CA29 $atom:12 $atom:13
106 bond:45CA28CA30 $atom:12 $atom:14
107 bond:423CA28CA36 $atom:12 $atom:15
108 bond:437CA28CA58 $atom:12 $atom:32
109 bond:451CA28CA63 $atom:12 $atom:33
110 bond:481CA28CA142 $atom:12 $atom:83
111 bond:46CA29CA30 $atom:13 $atom:14
112 bond:424CA29CA36 $atom:13 $atom:15
113 bond:438CA29CA58 $atom:13 $atom:32
114 bond:452CA29CA65 $atom:13 $atom:35
115 bond:482CA29CA142 $atom:13 $atom:83
116 bond:425CA30CA36 $atom:14 $atom:15
117 bond:439CA30CA56 $atom:14 $atom:30
118 bond:453CA30CA65 $atom:14 $atom:35
119 bond:483CA30CA142 $atom:14 $atom:83

120 bond:717CA30CA161 $atom:14 $atom:88
121 bond:718CA30CA162 $atom:14 $atom:89
122 bond:719CA30CA163 $atom:14 $atom:90
123 bond:720CA30CA164 $atom:14 $atom:91
124 bond:721CA30CA165 $atom:14 $atom:92
125 bond:722CA30CA166 $atom:14 $atom:93
126 bond:723CA30CA167 $atom:14 $atom:94
127 bond:724CA30CA168 $atom:14 $atom:95
128 bond:725CA30CA169 $atom:14 $atom:96
129 bond:726CA30CA170 $atom:14 $atom:97
130 bond:727CA30CA171 $atom:14 $atom:98
131 bond:728CA30CA172 $atom:14 $atom:99
132 bond:729CA30CA173 $atom:14 $atom:100
133 bond:730CA30CA174 $atom:14 $atom:101
134 bond:771CA30GC57 $atom:14 $atom:132
135 bond:47CA36CA37 $atom:15 $atom:16
136 bond:48CA36CA38 $atom:15 $atom:17
137 bond:49CA36CA39 $atom:15 $atom:18
138 bond:50CA36CA40 $atom:15 $atom:19
139 bond:484CA36CA56 $atom:15 $atom:30
140 bond:492CA36CA65 $atom:15 $atom:35
141 bond:500CA36CA104 $atom:15 $atom:57
142 bond:515CA36CA142 $atom:15 $atom:83
143 bond:731CA36CA161 $atom:15 $atom:88
144 bond:732CA36CA162 $atom:15 $atom:89
145 bond:733CA36CA163 $atom:15 $atom:90
146 bond:734CA36CA164 $atom:15 $atom:91
147 bond:735CA36CA165 $atom:15 $atom:92
148 bond:736CA36CA166 $atom:15 $atom:93
149 bond:737CA36CA167 $atom:15 $atom:94
150 bond:738CA36CA168 $atom:15 $atom:95
151 bond:739CA36CA169 $atom:15 $atom:96
152 bond:740CA36CA170 $atom:15 $atom:97
153 bond:741CA36CA171 $atom:15 $atom:98
154 bond:51CA37CA38 $atom:16 $atom:17
155 bond:52CA37CA39 $atom:16 $atom:18
156 bond:53CA37CA40 $atom:16 $atom:19
157 bond:54CA37CA41 $atom:16 $atom:20
158 bond:485CA37CA56 $atom:16 $atom:30
159 bond:493CA37CA69 $atom:16 $atom:39
160 bond:501CA37CA104 $atom:16 $atom:57
161 bond:508CA37CA118 $atom:16 $atom:65
162 bond:516CA37CA135 $atom:16 $atom:76
163 bond:742CA37CA172 $atom:16 $atom:99
164 bond:743CA37CA173 $atom:16 $atom:100
165 bond:744CA37CA174 $atom:16 $atom:101
166 bond:55CA38CA39 $atom:17 $atom:18
167 bond:56CA38CA40 $atom:17 $atom:19
168 bond:57CA38CA41 $atom:17 $atom:20
169 bond:58CA38CA42 $atom:17 $atom:21
170 bond:486CA38CA55 $atom:17 $atom:29

171 bond:494CA38CA69 $atom:17 $atom:39
172 bond:502CA38CA104 $atom:17 $atom:57
173 bond:509CA38CA118 $atom:17 $atom:65
174 bond:517CA38CA135 $atom:17 $atom:76
175 bond:773CA38GC57 $atom:17 $atom:132
176 bond:59CA39CA40 $atom:18 $atom:19
177 bond:60CA39CA41 $atom:18 $atom:20
178 bond:61CA39CA42 $atom:18 $atom:21
179 bond:62CA39CA43 $atom:18 $atom:22
180 bond:487CA39CA55 $atom:18 $atom:29
181 bond:495CA39CA66 $atom:18 $atom:36
182 bond:503CA39CA104 $atom:18 $atom:57
183 bond:510CA39CA118 $atom:18 $atom:65
184 bond:518CA39CA135 $atom:18 $atom:76
185 bond:63CA40CA41 $atom:19 $atom:20
186 bond:64CA40CA42 $atom:19 $atom:21
187 bond:65CA40CA43 $atom:19 $atom:22
188 bond:488CA40CA55 $atom:19 $atom:29
189 bond:496CA40CA69 $atom:19 $atom:39
190 bond:504CA40CA104 $atom:19 $atom:57
191 bond:511CA40CA118 $atom:19 $atom:65
192 bond:519CA40CA135 $atom:19 $atom:76
193 bond:66CA41CA42 $atom:20 $atom:21
194 bond:67CA41CA43 $atom:20 $atom:22
195 bond:489CA41CA52 $atom:20 $atom:26
196 bond:497CA41CA69 $atom:20 $atom:39
197 bond:505CA41CA104 $atom:20 $atom:57
198 bond:512CA41CA118 $atom:20 $atom:65
199 bond:520CA41CA131 $atom:20 $atom:72
200 bond:68CA42CA43 $atom:21 $atom:22
201 bond:490CA42CA52 $atom:21 $atom:26
202 bond:498CA42CA69 $atom:21 $atom:39
203 bond:506CA42CA104 $atom:21 $atom:57
204 bond:513CA42CA118 $atom:21 $atom:65
205 bond:521CA42CA131 $atom:21 $atom:72
206 bond:772CA42GC51 $atom:21 $atom:131
207 bond:491CA43CA55 $atom:22 $atom:29
208 bond:499CA43CA66 $atom:22 $atom:36
209 bond:507CA43CA104 $atom:22 $atom:57
210 bond:514CA43CA118 $atom:22 $atom:65
211 bond:522CA43CA131 $atom:22 $atom:72
212 bond:69CA49CA50 $atom:23 $atom:24
213 bond:70CA49CA51 $atom:23 $atom:25
214 bond:71CA49CA52 $atom:23 $atom:26
215 bond:72CA49CA53 $atom:23 $atom:27
216 bond:523CA49CA73 $atom:23 $atom:43
217 bond:533CA49CA104 $atom:23 $atom:57
218 bond:543CA49CA114 $atom:23 $atom:61
219 bond:553CA49CA127 $atom:23 $atom:68
220 bond:73CA50CA51 $atom:24 $atom:25
221 bond:74CA50CA52 $atom:24 $atom:26

222 bond:75CA50CA53 $atom:24 $atom:27
223 bond:76CA50CA54 $atom:24 $atom:28
224 bond:524CA50CA73 $atom:24 $atom:43
225 bond:534CA50CA104 $atom:24 $atom:57
226 bond:544CA50CA111 $atom:24 $atom:58
227 bond:554CA50CA127 $atom:24 $atom:68
228 bond:77CA51CA52 $atom:25 $atom:26
229 bond:78CA51CA53 $atom:25 $atom:27
230 bond:79CA51CA54 $atom:25 $atom:28
231 bond:80CA51CA55 $atom:25 $atom:29
232 bond:525CA51CA73 $atom:25 $atom:43
233 bond:535CA51CA104 $atom:25 $atom:57
234 bond:545CA51CA111 $atom:25 $atom:58
235 bond:555CA51CA127 $atom:25 $atom:68
236 bond:81CA52CA53 $atom:26 $atom:27
237 bond:82CA52CA54 $atom:26 $atom:28
238 bond:83CA52CA55 $atom:26 $atom:29
239 bond:84CA52CA56 $atom:26 $atom:30
240 bond:526CA52CA73 $atom:26 $atom:43
241 bond:536CA52CA104 $atom:26 $atom:57
242 bond:546CA52CA111 $atom:26 $atom:58
243 bond:556CA52CA131 $atom:26 $atom:72
244 bond:774CA52GC51 $atom:26 $atom:131
245 bond:85CA53CA54 $atom:27 $atom:28
246 bond:86CA53CA55 $atom:27 $atom:29
247 bond:87CA53CA56 $atom:27 $atom:30
248 bond:88CA53CA57 $atom:27 $atom:31
249 bond:527CA53CA70 $atom:27 $atom:40
250 bond:537CA53CA104 $atom:27 $atom:57
251 bond:547CA53CA111 $atom:27 $atom:58
252 bond:557CA53CA134 $atom:27 $atom:75
253 bond:89CA54CA55 $atom:28 $atom:29
254 bond:90CA54CA56 $atom:28 $atom:30
255 bond:91CA54CA57 $atom:28 $atom:31
256 bond:92CA54CA58 $atom:28 $atom:32
257 bond:528CA54CA66 $atom:28 $atom:36
258 bond:538CA54CA104 $atom:28 $atom:57
259 bond:548CA54CA111 $atom:28 $atom:58
260 bond:558CA54CA131 $atom:28 $atom:72
261 bond:93CA55CA56 $atom:29 $atom:30
262 bond:94CA55CA57 $atom:29 $atom:31
263 bond:95CA55CA58 $atom:29 $atom:32
264 bond:529CA55CA66 $atom:29 $atom:36
265 bond:539CA55CA104 $atom:29 $atom:57
266 bond:549CA55CA111 $atom:29 $atom:58
267 bond:559CA55CA134 $atom:29 $atom:75
268 bond:775CA55GC57 $atom:29 $atom:132
269 bond:96CA56CA57 $atom:30 $atom:31
270 bond:97CA56CA58 $atom:30 $atom:32
271 bond:530CA56CA66 $atom:30 $atom:36
272 bond:540CA56CA104 $atom:30 $atom:57

273 bond:550CA56CA111 $atom:30 $atom:58
274 bond:560CA56CA134 $atom:30 $atom:75
275 bond:745CA56CA170 $atom:30 $atom:97
276 bond:746CA56CA173 $atom:30 $atom:100
277 bond:747CA56CA174 $atom:30 $atom:101
278 bond:98CA57CA58 $atom:31 $atom:32
279 bond:531CA57CA66 $atom:31 $atom:36
280 bond:541CA57CA104 $atom:31 $atom:57
281 bond:551CA57CA111 $atom:31 $atom:58
282 bond:561CA57CA134 $atom:31 $atom:75
283 bond:532CA58CA66 $atom:32 $atom:36
284 bond:542CA58CA104 $atom:32 $atom:57
285 bond:552CA58CA111 $atom:32 $atom:58
286 bond:562CA58CA134 $atom:32 $atom:75
287 bond:99CA63CA64 $atom:33 $atom:34
288 bond:100CA63CA65 $atom:33 $atom:35
289 bond:101CA63CA66 $atom:33 $atom:36
290 bond:102CA63CA67 $atom:33 $atom:37
291 bond:563CA63CA102 $atom:33 $atom:55
292 bond:602CA63CA141 $atom:33 $atom:82
293 bond:103CA64CA65 $atom:34 $atom:35
294 bond:104CA64CA66 $atom:34 $atom:36
295 bond:105CA64CA67 $atom:34 $atom:37
296 bond:106CA64CA68 $atom:34 $atom:38
297 bond:564CA64CA102 $atom:34 $atom:55
298 bond:603CA64CA141 $atom:34 $atom:82
299 bond:107CA65CA66 $atom:35 $atom:36
300 bond:108CA65CA67 $atom:35 $atom:37
301 bond:109CA65CA68 $atom:35 $atom:38
302 bond:110CA65CA69 $atom:35 $atom:39
303 bond:565CA65CA102 $atom:35 $atom:55
304 bond:604CA65CA141 $atom:35 $atom:82
305 bond:748CA65CA167 $atom:35 $atom:94
306 bond:749CA65CA168 $atom:35 $atom:95
307 bond:750CA65CA169 $atom:35 $atom:96
308 bond:751CA65CA170 $atom:35 $atom:97
309 bond:752CA65CA171 $atom:35 $atom:98
310 bond:753CA65CA172 $atom:35 $atom:99
311 bond:754CA65CA173 $atom:35 $atom:100
312 bond:755CA65CA174 $atom:35 $atom:101
313 bond:776CA65GC57 $atom:35 $atom:132
314 bond:111CA66CA67 $atom:36 $atom:37
315 bond:112CA66CA68 $atom:36 $atom:38
316 bond:113CA66CA69 $atom:36 $atom:39
317 bond:114CA66CA70 $atom:36 $atom:40
318 bond:566CA66CA102 $atom:36 $atom:55
319 bond:584CA66CA111 $atom:36 $atom:58
320 bond:605CA66CA138 $atom:36 $atom:79
321 bond:115CA67CA68 $atom:37 $atom:38
322 bond:116CA67CA69 $atom:37 $atom:39
323 bond:117CA67CA70 $atom:37 $atom:40

324 bond:118CA67CA71 $atom:37 $atom:41
325 bond:567CA67CA102 $atom:37 $atom:55
326 bond:585CA67CA111 $atom:37 $atom:58
327 bond:606CA67CA137 $atom:37 $atom:78
328 bond:119CA68CA69 $atom:38 $atom:39
329 bond:120CA68CA70 $atom:38 $atom:40
330 bond:121CA68CA71 $atom:38 $atom:41
331 bond:122CA68CA72 $atom:38 $atom:42
332 bond:568CA68CA102 $atom:38 $atom:55
333 bond:586CA68CA114 $atom:38 $atom:61
334 bond:607CA68CA137 $atom:38 $atom:78
335 bond:123CA69CA70 $atom:39 $atom:40
336 bond:124CA69CA71 $atom:39 $atom:41
337 bond:125CA69CA72 $atom:39 $atom:42
338 bond:126CA69CA73 $atom:39 $atom:43
339 bond:569CA69CA102 $atom:39 $atom:55
340 bond:587CA69CA114 $atom:39 $atom:61
341 bond:608CA69CA137 $atom:39 $atom:78
342 bond:127CA70CA71 $atom:40 $atom:41
343 bond:128CA70CA72 $atom:40 $atom:42
344 bond:129CA70CA73 $atom:40 $atom:43
345 bond:130CA70CA74 $atom:40 $atom:44
346 bond:570CA70CA102 $atom:40 $atom:55
347 bond:588CA70CA114 $atom:40 $atom:61
348 bond:609CA70CA137 $atom:40 $atom:78
349 bond:131CA71CA72 $atom:41 $atom:42
350 bond:132CA71CA73 $atom:41 $atom:43
351 bond:133CA71CA74 $atom:41 $atom:44
352 bond:134CA71CA75 $atom:41 $atom:45
353 bond:571CA71CA102 $atom:41 $atom:55
354 bond:589CA71CA114 $atom:41 $atom:61
355 bond:610CA71CA137 $atom:41 $atom:78
356 bond:135CA72CA73 $atom:42 $atom:43
357 bond:136CA72CA74 $atom:42 $atom:44
358 bond:137CA72CA75 $atom:42 $atom:45
359 bond:138CA72CA76 $atom:42 $atom:46
360 bond:572CA72CA101 $atom:42 $atom:54
361 bond:590CA72CA114 $atom:42 $atom:61
362 bond:611CA72CA137 $atom:42 $atom:78
363 bond:139CA73CA74 $atom:43 $atom:44
364 bond:140CA73CA75 $atom:43 $atom:45
365 bond:141CA73CA76 $atom:43 $atom:46
366 bond:142CA73CA77 $atom:43 $atom:47
367 bond:573CA73CA101 $atom:43 $atom:54
368 bond:591CA73CA114 $atom:43 $atom:61
369 bond:612CA73CA133 $atom:43 $atom:74
370 bond:143CA74CA75 $atom:44 $atom:45
371 bond:144CA74CA76 $atom:44 $atom:46
372 bond:145CA74CA77 $atom:44 $atom:47
373 bond:146CA74CA78 $atom:44 $atom:48
374 bond:574CA74CA102 $atom:44 $atom:55

375 bond:592CA74CA114 $atom:44 $atom:61
376 bond:613CA74CA133 $atom:44 $atom:74
377 bond:147CA75CA76 $atom:45 $atom:46
378 bond:148CA75CA77 $atom:45 $atom:47
379 bond:149CA75CA78 $atom:45 $atom:48
380 bond:150CA75CA79 $atom:45 $atom:49
381 bond:575CA75CA101 $atom:45 $atom:54
382 bond:593CA75CA114 $atom:45 $atom:61
383 bond:614CA75CA133 $atom:45 $atom:74
384 bond:151CA76CA77 $atom:46 $atom:47
385 bond:152CA76CA78 $atom:46 $atom:48
386 bond:153CA76CA79 $atom:46 $atom:49
387 bond:154CA76CA80 $atom:46 $atom:50
388 bond:576CA76CA101 $atom:46 $atom:54
389 bond:594CA76CA114 $atom:46 $atom:61
390 bond:615CA76CA133 $atom:46 $atom:74
391 bond:155CA77CA78 $atom:47 $atom:48
392 bond:156CA77CA79 $atom:47 $atom:49
393 bond:157CA77CA80 $atom:47 $atom:50
394 bond:158CA77CA81 $atom:47 $atom:51
395 bond:577CA77CA101 $atom:47 $atom:54
396 bond:595CA77CA114 $atom:47 $atom:61
397 bond:616CA77CA133 $atom:47 $atom:74
398 bond:159CA78CA79 $atom:48 $atom:49
399 bond:160CA78CA80 $atom:48 $atom:50
400 bond:161CA78CA81 $atom:48 $atom:51
401 bond:162CA78CA82 $atom:48 $atom:52
402 bond:578CA78CA101 $atom:48 $atom:54
403 bond:596CA78CA114 $atom:48 $atom:61
404 bond:617CA78CA133 $atom:48 $atom:74
405 bond:163CA79CA80 $atom:49 $atom:50
406 bond:164CA79CA81 $atom:49 $atom:51
407 bond:165CA79CA82 $atom:49 $atom:52
408 bond:166CA79CA83 $atom:49 $atom:53
409 bond:579CA79CA101 $atom:49 $atom:54
410 bond:597CA79CA117 $atom:49 $atom:64
411 bond:618CA79CA133 $atom:49 $atom:74
412 bond:167CA80CA81 $atom:50 $atom:51
413 bond:168CA80CA82 $atom:50 $atom:52
414 bond:169CA80CA83 $atom:50 $atom:53
415 bond:580CA80CA101 $atom:50 $atom:54
416 bond:598CA80CA117 $atom:50 $atom:64
417 bond:619CA80CA129 $atom:50 $atom:70
418 bond:170CA81CA82 $atom:51 $atom:52
419 bond:171CA81CA83 $atom:51 $atom:53
420 bond:581CA81CA101 $atom:51 $atom:54
421 bond:599CA81CA117 $atom:51 $atom:64
422 bond:620CA81CA129 $atom:51 $atom:70
423 bond:172CA82CA83 $atom:52 $atom:53
424 bond:582CA82CA101 $atom:52 $atom:54
425 bond:600CA82CA117 $atom:52 $atom:64

426 bond:621CA82CA133 $atom:52 $atom:74
427 bond:583CA83CA101 $atom:53 $atom:54
428 bond:601CA83CA117 $atom:53 $atom:64
429 bond:622CA83CA129 $atom:53 $atom:70
430 bond:173CA101CA102 $atom:54 $atom:55
431 bond:174CA101CA103 $atom:54 $atom:56
432 bond:175CA101CA104 $atom:54 $atom:57
433 bond:623CA101CA114 $atom:54 $atom:61
434 bond:627CA101CA130 $atom:54 $atom:71
435 bond:176CA102CA103 $atom:55 $atom:56
436 bond:177CA102CA104 $atom:55 $atom:57
437 bond:624CA102CA114 $atom:55 $atom:61
438 bond:628CA102CA130 $atom:55 $atom:71
439 bond:178CA103CA104 $atom:56 $atom:57
440 bond:625CA103CA114 $atom:56 $atom:61
441 bond:629CA103CA126 $atom:56 $atom:67
442 bond:626CA104CA114 $atom:57 $atom:61
443 bond:630CA104CA126 $atom:57 $atom:67
444 bond:179CA111CA112 $atom:58 $atom:59
445 bond:180CA111CA113 $atom:58 $atom:60
446 bond:181CA111CA114 $atom:58 $atom:61
447 bond:182CA111CA115 $atom:58 $atom:62
448 bond:631CA111CA127 $atom:58 $atom:68
449 bond:183CA112CA113 $atom:59 $atom:60
450 bond:184CA112CA114 $atom:59 $atom:61
451 bond:185CA112CA115 $atom:59 $atom:62
452 bond:186CA112CA116 $atom:59 $atom:63
453 bond:632CA112CA126 $atom:59 $atom:67
454 bond:187CA113CA114 $atom:60 $atom:61
455 bond:188CA113CA115 $atom:60 $atom:62
456 bond:189CA113CA116 $atom:60 $atom:63
457 bond:190CA113CA117 $atom:60 $atom:64
458 bond:633CA113CA126 $atom:60 $atom:67
459 bond:191CA114CA115 $atom:61 $atom:62
460 bond:192CA114CA116 $atom:61 $atom:63
461 bond:193CA114CA117 $atom:61 $atom:64
462 bond:194CA114CA118 $atom:61 $atom:65
463 bond:634CA114CA126 $atom:61 $atom:67
464 bond:195CA115CA116 $atom:62 $atom:63
465 bond:196CA115CA117 $atom:62 $atom:64
466 bond:197CA115CA118 $atom:62 $atom:65
467 bond:198CA115CA119 $atom:62 $atom:66
468 bond:635CA115CA126 $atom:62 $atom:67
469 bond:199CA116CA117 $atom:63 $atom:64
470 bond:200CA116CA118 $atom:63 $atom:65
471 bond:201CA116CA119 $atom:63 $atom:66
472 bond:636CA116CA126 $atom:63 $atom:67
473 bond:202CA117CA118 $atom:64 $atom:65
474 bond:203CA117CA119 $atom:64 $atom:66
475 bond:637CA117CA126 $atom:64 $atom:67
476 bond:204CA118CA119 $atom:65 $atom:66

477 bond:638CA118CA126 $atom:65 $atom:67
478 bond:777CA118GC51 $atom:65 $atom:131
479 bond:639CA119CA126 $atom:66 $atom:67
480 bond:205CA126CA127 $atom:67 $atom:68
481 bond:206CA126CA128 $atom:67 $atom:69
482 bond:207CA126CA129 $atom:67 $atom:70
483 bond:208CA126CA130 $atom:67 $atom:71
484 bond:209CA127CA128 $atom:68 $atom:69
485 bond:210CA127CA129 $atom:68 $atom:70
486 bond:211CA127CA130 $atom:68 $atom:71
487 bond:212CA127CA131 $atom:68 $atom:72
488 bond:213CA128CA129 $atom:69 $atom:70
489 bond:214CA128CA130 $atom:69 $atom:71
490 bond:215CA128CA131 $atom:69 $atom:72
491 bond:216CA128CA132 $atom:69 $atom:73
492 bond:778CA128GC51 $atom:69 $atom:131
493 bond:217CA129CA130 $atom:70 $atom:71
494 bond:218CA129CA131 $atom:70 $atom:72
495 bond:219CA129CA132 $atom:70 $atom:73
496 bond:220CA129CA133 $atom:70 $atom:74
497 bond:221CA130CA131 $atom:71 $atom:72
498 bond:222CA130CA132 $atom:71 $atom:73
499 bond:223CA130CA133 $atom:71 $atom:74
500 bond:224CA130CA134 $atom:71 $atom:75
501 bond:225CA131CA132 $atom:72 $atom:73
502 bond:226CA131CA133 $atom:72 $atom:74
503 bond:227CA131CA134 $atom:72 $atom:75
504 bond:228CA131CA135 $atom:72 $atom:76
505 bond:229CA132CA133 $atom:73 $atom:74
506 bond:230CA132CA134 $atom:73 $atom:75
507 bond:231CA132CA135 $atom:73 $atom:76
508 bond:232CA132CA136 $atom:73 $atom:77
509 bond:233CA133CA134 $atom:74 $atom:75
510 bond:234CA133CA135 $atom:74 $atom:76
511 bond:235CA133CA136 $atom:74 $atom:77
512 bond:236CA133CA137 $atom:74 $atom:78
513 bond:237CA134CA135 $atom:75 $atom:76
514 bond:238CA134CA136 $atom:75 $atom:77
515 bond:239CA134CA137 $atom:75 $atom:78
516 bond:240CA134CA138 $atom:75 $atom:79
517 bond:241CA135CA136 $atom:76 $atom:77
518 bond:242CA135CA137 $atom:76 $atom:78
519 bond:243CA135CA138 $atom:76 $atom:79
520 bond:244CA135CA139 $atom:76 $atom:80
521 bond:245CA136CA137 $atom:77 $atom:78
522 bond:246CA136CA138 $atom:77 $atom:79
523 bond:247CA136CA139 $atom:77 $atom:80
524 bond:248CA136CA140 $atom:77 $atom:81
525 bond:249CA137CA138 $atom:78 $atom:79
526 bond:250CA137CA139 $atom:78 $atom:80
527 bond:251CA137CA140 $atom:78 $atom:81

528 bond:252CA137CA141 $atom:78 $atom:82
529 bond:253CA138CA139 $atom:79 $atom:80
530 bond:254CA138CA140 $atom:79 $atom:81
531 bond:255CA138CA141 $atom:79 $atom:82
532 bond:256CA138CA142 $atom:79 $atom:83
533 bond:257CA139CA140 $atom:80 $atom:81
534 bond:258CA139CA141 $atom:80 $atom:82
535 bond:259CA139CA142 $atom:80 $atom:83
536 bond:260CA139CA143 $atom:80 $atom:84
537 bond:779CA139GC57 $atom:80 $atom:132
538 bond:261CA140CA141 $atom:81 $atom:82
539 bond:262CA140CA142 $atom:81 $atom:83
540 bond:263CA140CA143 $atom:81 $atom:84
541 bond:264CA140CA144 $atom:81 $atom:85
542 bond:265CA141CA142 $atom:82 $atom:83
543 bond:266CA141CA143 $atom:82 $atom:84
544 bond:267CA141CA144 $atom:82 $atom:85
545 bond:268CA141CA145 $atom:82 $atom:86
546 bond:269CA142CA143 $atom:83 $atom:84
547 bond:270CA142CA144 $atom:83 $atom:85
548 bond:271CA142CA145 $atom:83 $atom:86
549 bond:272CA142CA146 $atom:83 $atom:87
550 bond:273CA143CA144 $atom:84 $atom:85
551 bond:274CA143CA145 $atom:84 $atom:86
552 bond:275CA143CA146 $atom:84 $atom:87
553 bond:767CA143CA172 $atom:84 $atom:99
554 bond:768CA143CA173 $atom:84 $atom:100
555 bond:769CA143CA174 $atom:84 $atom:101
556 bond:276CA144CA145 $atom:85 $atom:86
557 bond:277CA144CA146 $atom:85 $atom:87
558 bond:278CA145CA146 $atom:86 $atom:87
559 bond:756CA146CA161 $atom:87 $atom:88
560 bond:757CA146CA162 $atom:87 $atom:89
561 bond:758CA146CA163 $atom:87 $atom:90
562 bond:759CA146CA164 $atom:87 $atom:91
563 bond:760CA146CA165 $atom:87 $atom:92
564 bond:761CA146CA166 $atom:87 $atom:93
565 bond:762CA146CA167 $atom:87 $atom:94
566 bond:763CA146CA168 $atom:87 $atom:95
567 bond:764CA146CA169 $atom:87 $atom:96
568 bond:765CA146CA170 $atom:87 $atom:97
569 bond:766CA146CA171 $atom:87 $atom:98
570 bond:279CA161CA162 $atom:88 $atom:89
571 bond:280CA161CA163 $atom:88 $atom:90
572 bond:281CA161CA164 $atom:88 $atom:91
573 bond:282CA161CA165 $atom:88 $atom:92
574 bond:640CA161CA191 $atom:88 $atom:113
575 bond:654CA161CA198 $atom:88 $atom:117
576 bond:668CA161CA215 $atom:88 $atom:128
577 bond:784CA161GC204 $atom:88 $atom:134
578 bond:283CA162CA163 $atom:89 $atom:90

579 bond:284CA162CA164 $atom:89 $atom:91
580 bond:285CA162CA165 $atom:89 $atom:92
581 bond:286CA162CA166 $atom:89 $atom:93
582 bond:641CA162CA190 $atom:89 $atom:112
583 bond:655CA162CA198 $atom:89 $atom:117
584 bond:669CA162CA215 $atom:89 $atom:128
585 bond:287CA163CA164 $atom:90 $atom:91
586 bond:288CA163CA165 $atom:90 $atom:92
587 bond:289CA163CA166 $atom:90 $atom:93
588 bond:290CA163CA167 $atom:90 $atom:94
589 bond:642CA163CA190 $atom:90 $atom:112
590 bond:656CA163CA198 $atom:90 $atom:117
591 bond:670CA163CA215 $atom:90 $atom:128
592 bond:291CA164CA165 $atom:91 $atom:92
593 bond:292CA164CA166 $atom:91 $atom:93
594 bond:293CA164CA167 $atom:91 $atom:94
595 bond:294CA164CA168 $atom:91 $atom:95
596 bond:643CA164CA190 $atom:91 $atom:112
597 bond:657CA164CA198 $atom:91 $atom:117
598 bond:671CA164CA215 $atom:91 $atom:128
599 bond:295CA165CA166 $atom:92 $atom:93
600 bond:296CA165CA167 $atom:92 $atom:94
601 bond:297CA165CA168 $atom:92 $atom:95
602 bond:298CA165CA169 $atom:92 $atom:96
603 bond:644CA165CA190 $atom:92 $atom:112
604 bond:658CA165CA198 $atom:92 $atom:117
605 bond:672CA165CA215 $atom:92 $atom:128
606 bond:299CA166CA167 $atom:93 $atom:94
607 bond:300CA166CA168 $atom:93 $atom:95
608 bond:301CA166CA169 $atom:93 $atom:96
609 bond:302CA166CA170 $atom:93 $atom:97
610 bond:645CA166CA190 $atom:93 $atom:112
611 bond:659CA166CA198 $atom:93 $atom:117
612 bond:673CA166CA215 $atom:93 $atom:128
613 bond:780CA166GC63 $atom:93 $atom:133
614 bond:303CA167CA168 $atom:94 $atom:95
615 bond:304CA167CA169 $atom:94 $atom:96
616 bond:305CA167CA170 $atom:94 $atom:97
617 bond:306CA167CA171 $atom:94 $atom:98
618 bond:646CA167CA190 $atom:94 $atom:112
619 bond:660CA167CA198 $atom:94 $atom:117
620 bond:674CA167CA215 $atom:94 $atom:128
621 bond:307CA168CA169 $atom:95 $atom:96
622 bond:308CA168CA170 $atom:95 $atom:97
623 bond:309CA168CA171 $atom:95 $atom:98
624 bond:310CA168CA172 $atom:95 $atom:99
625 bond:647CA168CA190 $atom:95 $atom:112
626 bond:661CA168CA198 $atom:95 $atom:117
627 bond:675CA168CA211 $atom:95 $atom:124
628 bond:311CA169CA170 $atom:96 $atom:97
629 bond:312CA169CA171 $atom:96 $atom:98

| | | | |
|---|---|---|---|
| 630 | bond:313CA169CA172 | $atom:96 | $atom:99 |
| 631 | bond:314CA169CA173 | $atom:96 | $atom:100 |
| 632 | bond:648CA169CA186 | $atom:96 | $atom:108 |
| 633 | bond:662CA169CA198 | $atom:96 | $atom:117 |
| 634 | bond:676CA169CA211 | $atom:96 | $atom:124 |
| 635 | bond:315CA170CA171 | $atom:97 | $atom:98 |
| 636 | bond:316CA170CA172 | $atom:97 | $atom:99 |
| 637 | bond:317CA170CA173 | $atom:97 | $atom:100 |
| 638 | bond:318CA170CA174 | $atom:97 | $atom:101 |
| 639 | bond:649CA170CA186 | $atom:97 | $atom:108 |
| 640 | bond:663CA170CA198 | $atom:97 | $atom:117 |
| 641 | bond:677CA170CA211 | $atom:97 | $atom:124 |
| 642 | bond:319CA171CA172 | $atom:98 | $atom:99 |
| 643 | bond:320CA171CA173 | $atom:98 | $atom:100 |
| 644 | bond:321CA171CA174 | $atom:98 | $atom:101 |
| 645 | bond:650CA171CA186 | $atom:98 | $atom:108 |
| 646 | bond:664CA171CA198 | $atom:98 | $atom:117 |
| 647 | bond:678CA171CA211 | $atom:98 | $atom:124 |
| 648 | bond:322CA172CA173 | $atom:99 | $atom:100 |
| 649 | bond:323CA172CA174 | $atom:99 | $atom:101 |
| 650 | bond:651CA172CA182 | $atom:99 | $atom:104 |
| 651 | bond:665CA172CA198 | $atom:99 | $atom:117 |
| 652 | bond:679CA172CA211 | $atom:99 | $atom:124 |
| 653 | bond:324CA173CA174 | $atom:100 | $atom:101 |
| 654 | bond:652CA173CA182 | $atom:100 | $atom:104 |
| 655 | bond:666CA173CA198 | $atom:100 | $atom:117 |
| 656 | bond:680CA173CA211 | $atom:100 | $atom:124 |
| 657 | bond:653CA174CA182 | $atom:101 | $atom:104 |
| 658 | bond:667CA174CA198 | $atom:101 | $atom:117 |
| 659 | bond:681CA174CA211 | $atom:101 | $atom:124 |
| 660 | bond:325CA180CA181 | $atom:102 | $atom:103 |
| 661 | bond:326CA180CA182 | $atom:102 | $atom:104 |
| 662 | bond:327CA180CA183 | $atom:102 | $atom:105 |
| 663 | bond:328CA180CA184 | $atom:102 | $atom:106 |
| 664 | bond:682CA180CA203 | $atom:102 | $atom:122 |
| 665 | bond:695CA180CA211 | $atom:102 | $atom:124 |
| 666 | bond:801CA180CA172 | $atom:102 | $atom:233 |
| 667 | bond:788CA180CA180 | $atom:102 | $atom:236 |
| 668 | bond:329CA181CA182 | $atom:103 | $atom:104 |
| 669 | bond:330CA181CA183 | $atom:103 | $atom:105 |
| 670 | bond:331CA181CA184 | $atom:103 | $atom:106 |
| 671 | bond:332CA181CA185 | $atom:103 | $atom:107 |
| 672 | bond:683CA181CA199 | $atom:103 | $atom:118 |
| 673 | bond:696CA181CA211 | $atom:103 | $atom:124 |
| 674 | bond:802CA181CA172 | $atom:103 | $atom:233 |
| 675 | bond:789CA181CA181 | $atom:103 | $atom:237 |
| 676 | bond:333CA182CA183 | $atom:104 | $atom:105 |
| 677 | bond:334CA182CA184 | $atom:104 | $atom:106 |
| 678 | bond:335CA182CA185 | $atom:104 | $atom:107 |
| 679 | bond:336CA182CA186 | $atom:104 | $atom:108 |
| 680 | bond:684CA182CA202 | $atom:104 | $atom:121 |

681 bond:697CA182CA211 $atom:104 $atom:124
682 bond:803CA182CA172 $atom:104 $atom:233
683 bond:790CA182CA181 $atom:104 $atom:237
684 bond:337CA183CA184 $atom:105 $atom:106
685 bond:338CA183CA185 $atom:105 $atom:107
686 bond:339CA183CA186 $atom:105 $atom:108
687 bond:340CA183CA187 $atom:105 $atom:109
688 bond:685CA183CA202 $atom:105 $atom:121
689 bond:698CA183CA211 $atom:105 $atom:124
690 bond:804CA183CA172 $atom:105 $atom:233
691 bond:791CA183CA181 $atom:105 $atom:237
692 bond:341CA184CA185 $atom:106 $atom:107
693 bond:342CA184CA186 $atom:106 $atom:108
694 bond:343CA184CA187 $atom:106 $atom:109
695 bond:344CA184CA188 $atom:106 $atom:110
696 bond:686CA184CA203 $atom:106 $atom:122
697 bond:699CA184CA211 $atom:106 $atom:124
698 bond:805CA184CA172 $atom:106 $atom:233
699 bond:792CA184CA181 $atom:106 $atom:237
700 bond:345CA185CA186 $atom:107 $atom:108
701 bond:346CA185CA187 $atom:107 $atom:109
702 bond:347CA185CA188 $atom:107 $atom:110
703 bond:348CA185CA189 $atom:107 $atom:111
704 bond:349CA185CA190 $atom:107 $atom:112
705 bond:687CA185CA199 $atom:107 $atom:118
706 bond:700CA185CA211 $atom:107 $atom:124
707 bond:806CA185CA172 $atom:107 $atom:233
708 bond:793CA185CA181 $atom:107 $atom:237
709 bond:350CA186CA187 $atom:108 $atom:109
710 bond:351CA186CA188 $atom:108 $atom:110
711 bond:352CA186CA189 $atom:108 $atom:111
712 bond:353CA186CA190 $atom:108 $atom:112
713 bond:688CA186CA199 $atom:108 $atom:118
714 bond:701CA186CA211 $atom:108 $atom:124
715 bond:781CA186GC63 $atom:108 $atom:133
716 bond:807CA186CA172 $atom:108 $atom:233
717 bond:794CA186CA181 $atom:108 $atom:237
718 bond:354CA187CA188 $atom:109 $atom:110
719 bond:355CA187CA189 $atom:109 $atom:111
720 bond:356CA187CA190 $atom:109 $atom:112
721 bond:357CA187CA191 $atom:109 $atom:113
722 bond:689CA187CA202 $atom:109 $atom:121
723 bond:702CA187CA211 $atom:109 $atom:124
724 bond:808CA187CA172 $atom:109 $atom:233
725 bond:795CA187CA181 $atom:109 $atom:237
726 bond:358CA188CA189 $atom:110 $atom:111
727 bond:359CA188CA190 $atom:110 $atom:112
728 bond:360CA188CA191 $atom:110 $atom:113
729 bond:361CA188CA192 $atom:110 $atom:114
730 bond:690CA188CA199 $atom:110 $atom:118
731 bond:703CA188CA211 $atom:110 $atom:124

732 bond:809CA188CA172 $atom:110 $atom:233
733 bond:796CA188CA181 $atom:110 $atom:237
734 bond:362CA189CA190 $atom:111 $atom:112
735 bond:363CA189CA191 $atom:111 $atom:113
736 bond:364CA189CA192 $atom:111 $atom:114
737 bond:691CA189CA199 $atom:111 $atom:118
738 bond:704CA189CA211 $atom:111 $atom:124
739 bond:810CA189CA172 $atom:111 $atom:233
740 bond:797CA189CA185 $atom:111 $atom:241
741 bond:365CA190CA191 $atom:112 $atom:113
742 bond:366CA190CA192 $atom:112 $atom:114
743 bond:692CA190CA199 $atom:112 $atom:118
744 bond:705CA190CA211 $atom:112 $atom:124
745 bond:811CA190CA172 $atom:112 $atom:233
746 bond:798CA190CA185 $atom:112 $atom:241
747 bond:367CA191CA192 $atom:113 $atom:114
748 bond:693CA191CA199 $atom:113 $atom:118
749 bond:706CA191CA214 $atom:113 $atom:127
750 bond:785CA191GC204 $atom:113 $atom:134
751 bond:812CA191CA172 $atom:113 $atom:233
752 bond:799CA191CA189 $atom:113 $atom:245
753 bond:694CA192CA199 $atom:114 $atom:118
754 bond:707CA192CA214 $atom:114 $atom:127
755 bond:813CA192CA171 $atom:114 $atom:232
756 bond:800CA192CA189 $atom:114 $atom:245
757 bond:368CA196CA197 $atom:115 $atom:116
758 bond:369CA196CA198 $atom:115 $atom:117
759 bond:370CA196CA199 $atom:115 $atom:118
760 bond:371CA196CA200 $atom:115 $atom:119
761 bond:708CA196CA214 $atom:115 $atom:127
762 bond:372CA197CA198 $atom:116 $atom:117
763 bond:373CA197CA199 $atom:116 $atom:118
764 bond:374CA197CA200 $atom:116 $atom:119
765 bond:375CA197CA201 $atom:116 $atom:120
766 bond:709CA197CA217 $atom:116 $atom:130
767 bond:376CA198CA199 $atom:117 $atom:118
768 bond:377CA198CA200 $atom:117 $atom:119
769 bond:378CA198CA201 $atom:117 $atom:120
770 bond:379CA198CA202 $atom:117 $atom:121
771 bond:710CA198CA214 $atom:117 $atom:127
772 bond:782CA198GC63 $atom:117 $atom:133
773 bond:380CA199CA200 $atom:118 $atom:119
774 bond:381CA199CA201 $atom:118 $atom:120
775 bond:382CA199CA202 $atom:118 $atom:121
776 bond:383CA199CA203 $atom:118 $atom:122
777 bond:711CA199CA214 $atom:118 $atom:127
778 bond:384CA200CA201 $atom:119 $atom:120
779 bond:385CA200CA202 $atom:119 $atom:121
780 bond:386CA200CA203 $atom:119 $atom:122
781 bond:387CA200CA204 $atom:119 $atom:123
782 bond:712CA200CA214 $atom:119 $atom:127

| | | | |
|---|---|---|---|
| 783 | bond:388CA201CA202 | $atom:120 | $atom:121 |
| 784 | bond:389CA201CA203 | $atom:120 | $atom:122 |
| 785 | bond:390CA201CA204 | $atom:120 | $atom:123 |
| 786 | bond:713CA201CA217 | $atom:120 | $atom:130 |
| 787 | bond:391CA202CA203 | $atom:121 | $atom:122 |
| 788 | bond:392CA202CA204 | $atom:121 | $atom:123 |
| 789 | bond:714CA202CA214 | $atom:121 | $atom:127 |
| 790 | bond:393CA203CA204 | $atom:122 | $atom:123 |
| 791 | bond:715CA203CA214 | $atom:122 | $atom:127 |
| 792 | bond:716CA204CA214 | $atom:123 | $atom:127 |
| 793 | bond:786CA204GC204 | $atom:123 | $atom:134 |
| 794 | bond:394CA211CA212 | $atom:124 | $atom:125 |
| 795 | bond:395CA211CA213 | $atom:124 | $atom:126 |
| 796 | bond:396CA211CA214 | $atom:124 | $atom:127 |
| 797 | bond:397CA211CA215 | $atom:124 | $atom:128 |
| 798 | bond:398CA212CA213 | $atom:125 | $atom:126 |
| 799 | bond:399CA212CA214 | $atom:125 | $atom:127 |
| 800 | bond:400CA212CA215 | $atom:125 | $atom:128 |
| 801 | bond:401CA212CA216 | $atom:125 | $atom:129 |
| 802 | bond:402CA213CA214 | $atom:126 | $atom:127 |
| 803 | bond:403CA213CA215 | $atom:126 | $atom:128 |
| 804 | bond:404CA213CA216 | $atom:126 | $atom:129 |
| 805 | bond:405CA213CA217 | $atom:126 | $atom:130 |
| 806 | bond:406CA214CA215 | $atom:127 | $atom:128 |
| 807 | bond:407CA214CA216 | $atom:127 | $atom:129 |
| 808 | bond:408CA214CA217 | $atom:127 | $atom:130 |
| 809 | bond:409CA215CA216 | $atom:128 | $atom:129 |
| 810 | bond:410CA215CA217 | $atom:128 | $atom:130 |
| 811 | bond:783CA215GC63 | $atom:128 | $atom:133 |
| 812 | bond:411CA216CA217 | $atom:129 | $atom:130 |
| 813 | bond:787CA217GC204 | $atom:130 | $atom:134 |

**Table S2: CA bond coefficients data**

| **Type** | **no.** | ***K*** | ***$r_0$*** |
|---|---|---|---|
| bond_coeff | 1 | 10.0000 | 3.8080 |
| bond_coeff | 2 | 10.0000 | 5.5350 |
| bond_coeff | 3 | 10.0000 | 5.1060 |
| bond_coeff | 4 | 10.0000 | 6.0360 |
| bond_coeff | 5 | 10.0000 | 3.7910 |
| bond_coeff | 6 | 10.0000 | 5.5390 |
| bond_coeff | 7 | 10.0000 | 5.3180 |
| bond_coeff | 8 | 10.0000 | 6.0220 |
| bond_coeff | 9 | 10.0000 | 3.8150 |
| bond_coeff | 10 | 10.0000 | 5.4780 |
| bond_coeff | 11 | 10.0000 | 5.1960 |
| bond_coeff | 12 | 10.0000 | 6.2660 |
| bond_coeff | 13 | 10.0000 | 3.7860 |
| bond_coeff | 14 | 10.0000 | 5.6890 |
| bond_coeff | 15 | 10.0000 | 5.5750 |

```
bond_coeff   16   10.0000   6.4140
bond_coeff   17   10.0000   3.7970
bond_coeff   18   10.0000   5.4040
bond_coeff   19   10.0000   5.1450
bond_coeff   20   10.0000   6.0870
bond_coeff   21   10.0000   3.7820
bond_coeff   22   10.0000   5.6010
bond_coeff   23   10.0000   5.2550
bond_coeff   24   10.0000   6.1640
bond_coeff   25   10.0000   3.8010
bond_coeff   26   10.0000   5.4280
bond_coeff   27   10.0000   5.1820
bond_coeff   28   10.0000   6.5100
bond_coeff   29   10.0000   3.8390
bond_coeff   30   10.0000   5.5860
bond_coeff   31   10.0000   5.4640
bond_coeff   32   10.0000   6.3680
bond_coeff   33   10.0000   3.7890
bond_coeff   34   10.0000   5.3890
bond_coeff   35   10.0000   4.9630
bond_coeff   36   10.0000   5.9890
bond_coeff   37   10.0000   3.7970
bond_coeff   38   10.0000   5.4830
bond_coeff   39   10.0000   5.1970
bond_coeff   40   10.0000   5.7240
bond_coeff   41   10.0000   3.8010
bond_coeff   42   10.0000   5.4580
bond_coeff   43   10.0000   5.4520
bond_coeff   44   10.0000   3.7820
bond_coeff   45   10.0000   6.0410
bond_coeff   46   10.0000   3.7880
bond_coeff   47   10.0000   3.8160
bond_coeff   48   10.0000   5.8290
bond_coeff   49   10.0000   5.0540
bond_coeff   50   10.0000   6.1500
bond_coeff   51   10.0000   3.8260
bond_coeff   52   10.0000   5.1070
bond_coeff   53   10.0000   4.9270
bond_coeff   54   10.0000   6.1860
bond_coeff   55   10.0000   3.8150
bond_coeff   56   10.0000   5.5110
bond_coeff   57   10.0000   5.1040
bond_coeff   58   10.0000   6.2840
bond_coeff   59   10.0000   3.7930
bond_coeff   60   10.0000   5.4190
bond_coeff   61   10.0000   5.2220
bond_coeff   62   10.0000   6.1240
bond_coeff   63   10.0000   3.8020
bond_coeff   64   10.0000   5.4430
bond_coeff   65   10.0000   5.2060
bond_coeff   66   10.0000   3.7860
```

```
bond_coeff  67   10.0000  5.7160
bond_coeff  68   10.0000  3.8230
bond_coeff  69   10.0000  3.7780
bond_coeff  70   10.0000  5.4330
bond_coeff  71   10.0000  5.0010
bond_coeff  72   10.0000  6.2700
bond_coeff  73   10.0000  3.8400
bond_coeff  74   10.0000  5.4410
bond_coeff  75   10.0000  5.2620
bond_coeff  76   10.0000  6.3010
bond_coeff  77   10.0000  3.8040
bond_coeff  78   10.0000  5.4700
bond_coeff  79   10.0000  5.0000
bond_coeff  80   10.0000  6.3630
bond_coeff  81   10.0000  3.8250
bond_coeff  82   10.0000  5.3800
bond_coeff  83   10.0000  5.2550
bond_coeff  84   10.0000  6.3870
bond_coeff  85   10.0000  3.7900
bond_coeff  86   10.0000  5.4170
bond_coeff  87   10.0000  5.1150
bond_coeff  88   10.0000  5.8650
bond_coeff  89   10.0000  3.7840
bond_coeff  90   10.0000  5.5250
bond_coeff  91   10.0000  5.1770
bond_coeff  92   10.0000  6.1140
bond_coeff  93   10.0000  3.7940
bond_coeff  94   10.0000  5.5810
bond_coeff  95   10.0000  5.4890
bond_coeff  96   10.0000  3.8160
bond_coeff  97   10.0000  5.6170
bond_coeff  98   10.0000  3.8100
bond_coeff  99   10.0000  3.7910
bond_coeff  100  10.0000  5.3250
bond_coeff  101  10.0000  4.7380
bond_coeff  102  10.0000  5.8370
bond_coeff  103  10.0000  3.8170
bond_coeff  104  10.0000  5.4860
bond_coeff  105  10.0000  5.1030
bond_coeff  106  10.0000  6.3170
bond_coeff  107  10.0000  3.7990
bond_coeff  108  10.0000  5.2830
bond_coeff  109  10.0000  5.0850
bond_coeff  110  10.0000  6.0950
bond_coeff  111  10.0000  3.7790
bond_coeff  112  10.0000  5.5560
bond_coeff  113  10.0000  4.9920
bond_coeff  114  10.0000  6.4150
bond_coeff  115  10.0000  3.8030
bond_coeff  116  10.0000  5.2300
bond_coeff  117  10.0000  5.1560
```

| | | | |
|---|---|---|---|
| bond_coeff | 118 | 10.0000 | 6.6200 |
| bond_coeff | 119 | 10.0000 | 3.7940 |
| bond_coeff | 120 | 10.0000 | 5.4430 |
| bond_coeff | 121 | 10.0000 | 5.1390 |
| bond_coeff | 122 | 10.0000 | 6.4070 |
| bond_coeff | 123 | 10.0000 | 3.8090 |
| bond_coeff | 124 | 10.0000 | 5.3150 |
| bond_coeff | 125 | 10.0000 | 5.0910 |
| bond_coeff | 126 | 10.0000 | 6.1800 |
| bond_coeff | 127 | 10.0000 | 3.7950 |
| bond_coeff | 128 | 10.0000 | 5.5150 |
| bond_coeff | 129 | 10.0000 | 5.1040 |
| bond_coeff | 130 | 10.0000 | 6.1940 |
| bond_coeff | 131 | 10.0000 | 3.8080 |
| bond_coeff | 132 | 10.0000 | 5.4080 |
| bond_coeff | 133 | 10.0000 | 5.0790 |
| bond_coeff | 134 | 10.0000 | 6.3090 |
| bond_coeff | 135 | 10.0000 | 3.8190 |
| bond_coeff | 136 | 10.0000 | 5.4120 |
| bond_coeff | 137 | 10.0000 | 5.1370 |
| bond_coeff | 138 | 10.0000 | 6.3730 |
| bond_coeff | 139 | 10.0000 | 3.8070 |
| bond_coeff | 140 | 10.0000 | 5.4560 |
| bond_coeff | 141 | 10.0000 | 5.2120 |
| bond_coeff | 142 | 10.0000 | 6.1380 |
| bond_coeff | 143 | 10.0000 | 3.8020 |
| bond_coeff | 144 | 10.0000 | 5.4540 |
| bond_coeff | 145 | 10.0000 | 5.0370 |
| bond_coeff | 146 | 10.0000 | 6.1880 |
| bond_coeff | 147 | 10.0000 | 3.8040 |
| bond_coeff | 148 | 10.0000 | 5.4680 |
| bond_coeff | 149 | 10.0000 | 5.1930 |
| bond_coeff | 150 | 10.0000 | 6.1960 |
| bond_coeff | 151 | 10.0000 | 3.7930 |
| bond_coeff | 152 | 10.0000 | 5.3860 |
| bond_coeff | 153 | 10.0000 | 4.9280 |
| bond_coeff | 154 | 10.0000 | 6.0480 |
| bond_coeff | 155 | 10.0000 | 3.8230 |
| bond_coeff | 156 | 10.0000 | 5.3960 |
| bond_coeff | 157 | 10.0000 | 4.9720 |
| bond_coeff | 158 | 10.0000 | 6.2650 |
| bond_coeff | 159 | 10.0000 | 3.7780 |
| bond_coeff | 160 | 10.0000 | 5.3010 |
| bond_coeff | 161 | 10.0000 | 5.0670 |
| bond_coeff | 162 | 10.0000 | 6.1780 |
| bond_coeff | 163 | 10.0000 | 3.7780 |
| bond_coeff | 164 | 10.0000 | 5.4560 |
| bond_coeff | 165 | 10.0000 | 5.0310 |
| bond_coeff | 166 | 10.0000 | 6.3990 |
| bond_coeff | 167 | 10.0000 | 3.8000 |
| bond_coeff | 168 | 10.0000 | 5.3480 |

```
bond_coeff  169  10.0000  5.2020
bond_coeff  170  10.0000  3.7870
bond_coeff  171  10.0000  5.4230
bond_coeff  172  10.0000  3.8250
bond_coeff  173  10.0000  3.7960
bond_coeff  174  10.0000  5.2710
bond_coeff  175  10.0000  5.3160
bond_coeff  176  10.0000  3.7880
bond_coeff  177  10.0000  5.3950
bond_coeff  178  10.0000  3.8010
bond_coeff  179  10.0000  3.8220
bond_coeff  180  10.0000  5.5080
bond_coeff  181  10.0000  4.9800
bond_coeff  182  10.0000  6.0400
bond_coeff  183  10.0000  3.8030
bond_coeff  184  10.0000  5.4260
bond_coeff  185  10.0000  5.0960
bond_coeff  186  10.0000  5.9170
bond_coeff  187  10.0000  3.7940
bond_coeff  188  10.0000  5.3600
bond_coeff  189  10.0000  4.6610
bond_coeff  190  10.0000  6.1790
bond_coeff  191  10.0000  3.7990
bond_coeff  192  10.0000  5.2730
bond_coeff  193  10.0000  5.2790
bond_coeff  194  10.0000  6.1850
bond_coeff  195  10.0000  3.7740
bond_coeff  196  10.0000  5.4800
bond_coeff  197  10.0000  5.1100
bond_coeff  198  10.0000  5.9550
bond_coeff  199  10.0000  3.7970
bond_coeff  200  10.0000  5.6360
bond_coeff  201  10.0000  5.5880
bond_coeff  202  10.0000  3.7910
bond_coeff  203  10.0000  5.6460
bond_coeff  204  10.0000  3.8050
bond_coeff  205  10.0000  3.7860
bond_coeff  206  10.0000  5.5370
bond_coeff  207  10.0000  5.0510
bond_coeff  208  10.0000  5.9930
bond_coeff  209  10.0000  3.8120
bond_coeff  210  10.0000  5.5070
bond_coeff  211  10.0000  5.2970
bond_coeff  212  10.0000  6.0660
bond_coeff  213  10.0000  3.8060
bond_coeff  214  10.0000  5.4860
bond_coeff  215  10.0000  4.9220
bond_coeff  216  10.0000  6.2320
bond_coeff  217  10.0000  3.7910
bond_coeff  218  10.0000  5.3760
bond_coeff  219  10.0000  5.1810
```

```
bond_coeff 220 10.0000 6.7100
bond_coeff 221 10.0000 3.8070
bond_coeff 222 10.0000 5.3360
bond_coeff 223 10.0000 5.2090
bond_coeff 224 10.0000 6.2870
bond_coeff 225 10.0000 3.8030
bond_coeff 226 10.0000 5.4570
bond_coeff 227 10.0000 5.0650
bond_coeff 228 10.0000 6.1440
bond_coeff 229 10.0000 3.8260
bond_coeff 230 10.0000 5.5000
bond_coeff 231 10.0000 5.2380
bond_coeff 232 10.0000 6.3190
bond_coeff 233 10.0000 3.7880
bond_coeff 234 10.0000 5.4500
bond_coeff 235 10.0000 5.0960
bond_coeff 236 10.0000 6.4330
bond_coeff 237 10.0000 3.8110
bond_coeff 238 10.0000 5.4290
bond_coeff 239 10.0000 5.1700
bond_coeff 240 10.0000 6.1460
bond_coeff 241 10.0000 3.8370
bond_coeff 242 10.0000 5.3190
bond_coeff 243 10.0000 4.8140
bond_coeff 244 10.0000 5.9860
bond_coeff 245 10.0000 3.7970
bond_coeff 246 10.0000 5.4910
bond_coeff 247 10.0000 5.2230
bond_coeff 248 10.0000 6.5580
bond_coeff 249 10.0000 3.8000
bond_coeff 250 10.0000 5.4320
bond_coeff 251 10.0000 5.2500
bond_coeff 252 10.0000 6.5010
bond_coeff 253 10.0000 3.8080
bond_coeff 254 10.0000 5.3880
bond_coeff 255 10.0000 5.0410
bond_coeff 256 10.0000 6.2360
bond_coeff 257 10.0000 3.8110
bond_coeff 258 10.0000 5.3680
bond_coeff 259 10.0000 5.0670
bond_coeff 260 10.0000 6.3060
bond_coeff 261 10.0000 3.7950
bond_coeff 262 10.0000 5.5020
bond_coeff 263 10.0000 5.2840
bond_coeff 264 10.0000 6.2650
bond_coeff 265 10.0000 3.8270
bond_coeff 266 10.0000 5.4820
bond_coeff 267 10.0000 5.0960
bond_coeff 268 10.0000 6.0740
bond_coeff 269 10.0000 3.8180
bond_coeff 270 10.0000 5.4600
```

| | | | |
|---|---|---|---|
| bond_coeff | 271 | 10.0000 | 5.2570 |
| bond_coeff | 272 | 10.0000 | 6.2310 |
| bond_coeff | 273 | 10.0000 | 3.8060 |
| bond_coeff | 274 | 10.0000 | 5.6310 |
| bond_coeff | 275 | 10.0000 | 5.3260 |
| bond_coeff | 276 | 10.0000 | 3.7920 |
| bond_coeff | 277 | 10.0000 | 5.4020 |
| bond_coeff | 278 | 10.0000 | 3.7920 |
| bond_coeff | 279 | 10.0000 | 3.7940 |
| bond_coeff | 280 | 10.0000 | 5.1890 |
| bond_coeff | 281 | 10.0000 | 4.7980 |
| bond_coeff | 282 | 10.0000 | 6.0650 |
| bond_coeff | 283 | 10.0000 | 3.8000 |
| bond_coeff | 284 | 10.0000 | 5.4520 |
| bond_coeff | 285 | 10.0000 | 5.1840 |
| bond_coeff | 286 | 10.0000 | 6.0380 |
| bond_coeff | 287 | 10.0000 | 3.7900 |
| bond_coeff | 288 | 10.0000 | 5.4570 |
| bond_coeff | 289 | 10.0000 | 5.0160 |
| bond_coeff | 290 | 10.0000 | 5.9550 |
| bond_coeff | 291 | 10.0000 | 3.7890 |
| bond_coeff | 292 | 10.0000 | 5.4550 |
| bond_coeff | 293 | 10.0000 | 5.0340 |
| bond_coeff | 294 | 10.0000 | 5.7330 |
| bond_coeff | 295 | 10.0000 | 3.7970 |
| bond_coeff | 296 | 10.0000 | 5.3570 |
| bond_coeff | 297 | 10.0000 | 4.8360 |
| bond_coeff | 298 | 10.0000 | 5.7900 |
| bond_coeff | 299 | 10.0000 | 3.8040 |
| bond_coeff | 300 | 10.0000 | 5.5300 |
| bond_coeff | 301 | 10.0000 | 5.1600 |
| bond_coeff | 302 | 10.0000 | 5.8420 |
| bond_coeff | 303 | 10.0000 | 3.7950 |
| bond_coeff | 304 | 10.0000 | 5.3450 |
| bond_coeff | 305 | 10.0000 | 4.6470 |
| bond_coeff | 306 | 10.0000 | 6.0390 |
| bond_coeff | 307 | 10.0000 | 3.7650 |
| bond_coeff | 308 | 10.0000 | 5.2950 |
| bond_coeff | 309 | 10.0000 | 5.1030 |
| bond_coeff | 310 | 10.0000 | 5.9760 |
| bond_coeff | 311 | 10.0000 | 3.7940 |
| bond_coeff | 312 | 10.0000 | 5.3520 |
| bond_coeff | 313 | 10.0000 | 4.8620 |
| bond_coeff | 314 | 10.0000 | 5.9260 |
| bond_coeff | 315 | 10.0000 | 3.8110 |
| bond_coeff | 316 | 10.0000 | 5.5680 |
| bond_coeff | 317 | 10.0000 | 5.2640 |
| bond_coeff | 318 | 10.0000 | 6.6950 |
| bond_coeff | 319 | 10.0000 | 3.7760 |
| bond_coeff | 320 | 10.0000 | 5.3010 |
| bond_coeff | 321 | 10.0000 | 4.9830 |

| | | | |
|---|---|---|---|
| bond_coeff | 322 | 10.0000 | 3.7850 |
| bond_coeff | 323 | 10.0000 | 5.0810 |
| bond_coeff | 324 | 10.0000 | 3.7880 |
| bond_coeff | 325 | 10.0000 | 3.7970 |
| bond_coeff | 326 | 10.0000 | 5.4190 |
| bond_coeff | 327 | 10.0000 | 5.3550 |
| bond_coeff | 328 | 10.0000 | 6.5460 |
| bond_coeff | 329 | 10.0000 | 3.8050 |
| bond_coeff | 330 | 10.0000 | 5.4350 |
| bond_coeff | 331 | 10.0000 | 5.0840 |
| bond_coeff | 332 | 10.0000 | 5.3980 |
| bond_coeff | 333 | 10.0000 | 3.7970 |
| bond_coeff | 334 | 10.0000 | 5.4050 |
| bond_coeff | 335 | 10.0000 | 5.0020 |
| bond_coeff | 336 | 10.0000 | 5.0160 |
| bond_coeff | 337 | 10.0000 | 3.8000 |
| bond_coeff | 338 | 10.0000 | 5.8510 |
| bond_coeff | 339 | 10.0000 | 5.4730 |
| bond_coeff | 340 | 10.0000 | 6.0090 |
| bond_coeff | 341 | 10.0000 | 3.8030 |
| bond_coeff | 342 | 10.0000 | 5.5860 |
| bond_coeff | 343 | 10.0000 | 5.0090 |
| bond_coeff | 344 | 10.0000 | 5.8940 |
| bond_coeff | 345 | 10.0000 | 3.8010 |
| bond_coeff | 346 | 10.0000 | 5.1180 |
| bond_coeff | 347 | 10.0000 | 4.9180 |
| bond_coeff | 348 | 10.0000 | 5.2250 |
| bond_coeff | 349 | 10.0000 | 6.5420 |
| bond_coeff | 350 | 10.0000 | 3.8040 |
| bond_coeff | 351 | 10.0000 | 5.8300 |
| bond_coeff | 352 | 10.0000 | 6.0770 |
| bond_coeff | 353 | 10.0000 | 4.8820 |
| bond_coeff | 354 | 10.0000 | 3.8180 |
| bond_coeff | 355 | 10.0000 | 6.2570 |
| bond_coeff | 356 | 10.0000 | 5.4990 |
| bond_coeff | 357 | 10.0000 | 6.9520 |
| bond_coeff | 358 | 10.0000 | 3.7830 |
| bond_coeff | 359 | 10.0000 | 5.3280 |
| bond_coeff | 360 | 10.0000 | 6.0360 |
| bond_coeff | 361 | 10.0000 | 6.3240 |
| bond_coeff | 362 | 10.0000 | 3.7790 |
| bond_coeff | 363 | 10.0000 | 5.4800 |
| bond_coeff | 364 | 10.0000 | 4.6370 |
| bond_coeff | 365 | 10.0000 | 3.8040 |
| bond_coeff | 366 | 10.0000 | 5.5830 |
| bond_coeff | 367 | 10.0000 | 3.7980 |
| bond_coeff | 368 | 10.0000 | 3.8270 |
| bond_coeff | 369 | 10.0000 | 6.0790 |
| bond_coeff | 370 | 10.0000 | 5.7620 |
| bond_coeff | 371 | 10.0000 | 6.9660 |
| bond_coeff | 372 | 10.0000 | 3.7990 |

```
bond_coeff 373 10.0000 5.4410
bond_coeff 374 10.0000 5.6300
bond_coeff 375 10.0000 6.7520
bond_coeff 376 10.0000 3.7920
bond_coeff 377 10.0000 5.4770
bond_coeff 378 10.0000 5.3220
bond_coeff 379 10.0000 5.5450
bond_coeff 380 10.0000 3.8000
bond_coeff 381 10.0000 5.6900
bond_coeff 382 10.0000 5.2210
bond_coeff 383 10.0000 5.9220
bond_coeff 384 10.0000 3.7910
bond_coeff 385 10.0000 5.6170
bond_coeff 386 10.0000 5.2970
bond_coeff 387 10.0000 6.6250
bond_coeff 388 10.0000 3.7830
bond_coeff 389 10.0000 5.2610
bond_coeff 390 10.0000 5.1920
bond_coeff 391 10.0000 3.7920
bond_coeff 392 10.0000 5.5160
bond_coeff 393 10.0000 3.7990
bond_coeff 394 10.0000 3.7940
bond_coeff 395 10.0000 5.2230
bond_coeff 396 10.0000 4.8160
bond_coeff 397 10.0000 5.9840
bond_coeff 398 10.0000 3.8010
bond_coeff 399 10.0000 5.4940
bond_coeff 400 10.0000 5.4280
bond_coeff 401 10.0000 5.7790
bond_coeff 402 10.0000 3.8070
bond_coeff 403 10.0000 5.7290
bond_coeff 404 10.0000 5.3240
bond_coeff 405 10.0000 5.5120
bond_coeff 406 10.0000 3.8070
bond_coeff 407 10.0000 5.6260
bond_coeff 408 10.0000 5.2360
bond_coeff 409 10.0000 3.7670
bond_coeff 410 10.0000 5.5210
bond_coeff 411 10.0000 3.7740
bond_coeff 412 10.0000 11.7320
bond_coeff 413 10.0000 9.6780
bond_coeff 414 10.0000 6.2760
bond_coeff 415 10.0000 8.6050
bond_coeff 416 10.0000 10.2910
bond_coeff 417 10.0000 7.9840
bond_coeff 418 10.0000 7.0200
bond_coeff 419 10.0000 10.1870
bond_coeff 420 10.0000 10.6300
bond_coeff 421 10.0000 6.9130
bond_coeff 422 10.0000 7.3520
bond_coeff 423 10.0000 10.5840
```

| | | | |
|---|---|---|---|
| bond_coeff | 424 | 10.0000 | 9.7110 |
| bond_coeff | 425 | 10.0000 | 6.9140 |
| bond_coeff | 426 | 10.0000 | 11.3420 |
| bond_coeff | 427 | 10.0000 | 11.9770 |
| bond_coeff | 428 | 10.0000 | 8.8970 |
| bond_coeff | 429 | 10.0000 | 6.7170 |
| bond_coeff | 430 | 10.0000 | 9.2980 |
| bond_coeff | 431 | 10.0000 | 10.1150 |
| bond_coeff | 432 | 10.0000 | 7.1040 |
| bond_coeff | 433 | 10.0000 | 6.8230 |
| bond_coeff | 434 | 10.0000 | 10.5250 |
| bond_coeff | 435 | 10.0000 | 11.4810 |
| bond_coeff | 436 | 10.0000 | 10.1550 |
| bond_coeff | 437 | 10.0000 | 11.2390 |
| bond_coeff | 438 | 10.0000 | 14.6020 |
| bond_coeff | 439 | 10.0000 | 15.1890 |
| bond_coeff | 440 | 10.0000 | 21.8860 |
| bond_coeff | 441 | 10.0000 | 22.1930 |
| bond_coeff | 442 | 10.0000 | 19.3760 |
| bond_coeff | 443 | 10.0000 | 17.0070 |
| bond_coeff | 444 | 10.0000 | 18.0260 |
| bond_coeff | 445 | 10.0000 | 18.1140 |
| bond_coeff | 446 | 10.0000 | 14.4880 |
| bond_coeff | 447 | 10.0000 | 13.4330 |
| bond_coeff | 448 | 10.0000 | 15.9670 |
| bond_coeff | 449 | 10.0000 | 15.1170 |
| bond_coeff | 450 | 10.0000 | 11.5700 |
| bond_coeff | 451 | 10.0000 | 12.7180 |
| bond_coeff | 452 | 10.0000 | 15.5470 |
| bond_coeff | 453 | 10.0000 | 13.9290 |
| bond_coeff | 454 | 10.0000 | 24.3040 |
| bond_coeff | 455 | 10.0000 | 21.7030 |
| bond_coeff | 456 | 10.0000 | 21.1200 |
| bond_coeff | 457 | 10.0000 | 24.5410 |
| bond_coeff | 458 | 10.0000 | 24.5210 |
| bond_coeff | 459 | 10.0000 | 21.5570 |
| bond_coeff | 460 | 10.0000 | 23.1150 |
| bond_coeff | 461 | 10.0000 | 24.1200 |
| bond_coeff | 462 | 10.0000 | 19.1830 |
| bond_coeff | 463 | 10.0000 | 21.2580 |
| bond_coeff | 464 | 10.0000 | 18.9230 |
| bond_coeff | 465 | 10.0000 | 18.2020 |
| bond_coeff | 466 | 10.0000 | 21.9520 |
| bond_coeff | 467 | 10.0000 | 23.1800 |
| bond_coeff | 468 | 10.0000 | 21.4120 |
| bond_coeff | 469 | 10.0000 | 22.6350 |
| bond_coeff | 470 | 10.0000 | 19.8710 |
| bond_coeff | 471 | 10.0000 | 19.8260 |
| bond_coeff | 472 | 10.0000 | 16.0410 |
| bond_coeff | 473 | 10.0000 | 16.3390 |
| bond_coeff | 474 | 10.0000 | 19.0680 |

| | | | |
|---|---|---|---|
| bond_coeff | 475 | 10.0000 | 17.5210 |
| bond_coeff | 476 | 10.0000 | 14.0260 |
| bond_coeff | 477 | 10.0000 | 15.1130 |
| bond_coeff | 478 | 10.0000 | 16.7200 |
| bond_coeff | 479 | 10.0000 | 13.9580 |
| bond_coeff | 480 | 10.0000 | 11.9240 |
| bond_coeff | 481 | 10.0000 | 14.7080 |
| bond_coeff | 482 | 10.0000 | 14.5860 |
| bond_coeff | 483 | 10.0000 | 11.1610 |
| bond_coeff | 484 | 10.0000 | 12.8750 |
| bond_coeff | 485 | 10.0000 | 12.4480 |
| bond_coeff | 486 | 10.0000 | 14.4690 |
| bond_coeff | 487 | 10.0000 | 12.1620 |
| bond_coeff | 488 | 10.0000 | 8.9750 |
| bond_coeff | 489 | 10.0000 | 10.1130 |
| bond_coeff | 490 | 10.0000 | 11.8140 |
| bond_coeff | 491 | 10.0000 | 9.2420 |
| bond_coeff | 492 | 10.0000 | 13.6720 |
| bond_coeff | 493 | 10.0000 | 13.4410 |
| bond_coeff | 494 | 10.0000 | 16.8730 |
| bond_coeff | 495 | 10.0000 | 17.0960 |
| bond_coeff | 496 | 10.0000 | 14.1370 |
| bond_coeff | 497 | 10.0000 | 15.4990 |
| bond_coeff | 498 | 10.0000 | 18.8870 |
| bond_coeff | 499 | 10.0000 | 18.0160 |
| bond_coeff | 500 | 10.0000 | 23.0610 |
| bond_coeff | 501 | 10.0000 | 20.1930 |
| bond_coeff | 502 | 10.0000 | 21.3220 |
| bond_coeff | 503 | 10.0000 | 21.7040 |
| bond_coeff | 504 | 10.0000 | 18.1920 |
| bond_coeff | 505 | 10.0000 | 16.9170 |
| bond_coeff | 506 | 10.0000 | 19.3710 |
| bond_coeff | 507 | 10.0000 | 18.9470 |
| bond_coeff | 508 | 10.0000 | 23.0140 |
| bond_coeff | 509 | 10.0000 | 22.3430 |
| bond_coeff | 510 | 10.0000 | 22.4550 |
| bond_coeff | 511 | 10.0000 | 19.8990 |
| bond_coeff | 512 | 10.0000 | 17.4770 |
| bond_coeff | 513 | 10.0000 | 18.0740 |
| bond_coeff | 514 | 10.0000 | 18.0840 |
| bond_coeff | 515 | 10.0000 | 9.0320 |
| bond_coeff | 516 | 10.0000 | 7.1680 |
| bond_coeff | 517 | 10.0000 | 9.3450 |
| bond_coeff | 518 | 10.0000 | 11.0630 |
| bond_coeff | 519 | 10.0000 | 8.5590 |
| bond_coeff | 520 | 10.0000 | 7.7400 |
| bond_coeff | 521 | 10.0000 | 10.8390 |
| bond_coeff | 522 | 10.0000 | 11.3780 |
| bond_coeff | 523 | 10.0000 | 12.0000 |
| bond_coeff | 524 | 10.0000 | 14.1820 |
| bond_coeff | 525 | 10.0000 | 15.3760 |

| | | | |
|---|---|---|---|
| bond_coeff | 526 | 10.0000 | 12.0650 |
| bond_coeff | 527 | 10.0000 | 10.5920 |
| bond_coeff | 528 | 10.0000 | 12.6810 |
| bond_coeff | 529 | 10.0000 | 11.4260 |
| bond_coeff | 530 | 10.0000 | 7.6940 |
| bond_coeff | 531 | 10.0000 | 8.4830 |
| bond_coeff | 532 | 10.0000 | 10.7280 |
| bond_coeff | 533 | 10.0000 | 6.4850 |
| bond_coeff | 534 | 10.0000 | 9.3990 |
| bond_coeff | 535 | 10.0000 | 11.8980 |
| bond_coeff | 536 | 10.0000 | 10.6270 |
| bond_coeff | 537 | 10.0000 | 10.9750 |
| bond_coeff | 538 | 10.0000 | 14.2450 |
| bond_coeff | 539 | 10.0000 | 15.5560 |
| bond_coeff | 540 | 10.0000 | 15.0200 |
| bond_coeff | 541 | 10.0000 | 16.4180 |
| bond_coeff | 542 | 10.0000 | 19.4150 |
| bond_coeff | 543 | 10.0000 | 7.2100 |
| bond_coeff | 544 | 10.0000 | 6.7620 |
| bond_coeff | 545 | 10.0000 | 9.4630 |
| bond_coeff | 546 | 10.0000 | 11.9000 |
| bond_coeff | 547 | 10.0000 | 11.8870 |
| bond_coeff | 548 | 10.0000 | 12.4480 |
| bond_coeff | 549 | 10.0000 | 15.3400 |
| bond_coeff | 550 | 10.0000 | 16.8300 |
| bond_coeff | 551 | 10.0000 | 16.6920 |
| bond_coeff | 552 | 10.0000 | 18.3780 |
| bond_coeff | 553 | 10.0000 | 5.1390 |
| bond_coeff | 554 | 10.0000 | 8.5300 |
| bond_coeff | 555 | 10.0000 | 8.6670 |
| bond_coeff | 556 | 10.0000 | 7.3460 |
| bond_coeff | 557 | 10.0000 | 9.7760 |
| bond_coeff | 558 | 10.0000 | 12.5950 |
| bond_coeff | 559 | 10.0000 | 11.1530 |
| bond_coeff | 560 | 10.0000 | 9.3130 |
| bond_coeff | 561 | 10.0000 | 12.5740 |
| bond_coeff | 562 | 10.0000 | 14.9140 |
| bond_coeff | 563 | 10.0000 | 21.1280 |
| bond_coeff | 564 | 10.0000 | 21.5940 |
| bond_coeff | 565 | 10.0000 | 20.1470 |
| bond_coeff | 566 | 10.0000 | 17.2960 |
| bond_coeff | 567 | 10.0000 | 16.5650 |
| bond_coeff | 568 | 10.0000 | 16.8650 |
| bond_coeff | 569 | 10.0000 | 14.4820 |
| bond_coeff | 570 | 10.0000 | 11.6910 |
| bond_coeff | 571 | 10.0000 | 12.6460 |
| bond_coeff | 572 | 10.0000 | 12.3310 |
| bond_coeff | 573 | 10.0000 | 9.1530 |
| bond_coeff | 574 | 10.0000 | 8.1370 |
| bond_coeff | 575 | 10.0000 | 9.4730 |
| bond_coeff | 576 | 10.0000 | 7.6160 |

| | | | |
|---|---|---|---|
| bond_coeff | 577 | 10.0000 | 4.0810 |
| bond_coeff | 578 | 10.0000 | 6.3750 |
| bond_coeff | 579 | 10.0000 | 8.0220 |
| bond_coeff | 580 | 10.0000 | 5.6870 |
| bond_coeff | 581 | 10.0000 | 5.6650 |
| bond_coeff | 582 | 10.0000 | 9.2420 |
| bond_coeff | 583 | 10.0000 | 10.0830 |
| bond_coeff | 584 | 10.0000 | 22.7080 |
| bond_coeff | 585 | 10.0000 | 23.8600 |
| bond_coeff | 586 | 10.0000 | 24.7520 |
| bond_coeff | 587 | 10.0000 | 21.3270 |
| bond_coeff | 588 | 10.0000 | 19.7640 |
| bond_coeff | 589 | 10.0000 | 22.0460 |
| bond_coeff | 590 | 10.0000 | 21.1140 |
| bond_coeff | 591 | 10.0000 | 17.3630 |
| bond_coeff | 592 | 10.0000 | 18.4300 |
| bond_coeff | 593 | 10.0000 | 20.6670 |
| bond_coeff | 594 | 10.0000 | 18.1110 |
| bond_coeff | 595 | 10.0000 | 15.7280 |
| bond_coeff | 596 | 10.0000 | 18.9020 |
| bond_coeff | 597 | 10.0000 | 19.7900 |
| bond_coeff | 598 | 10.0000 | 16.1500 |
| bond_coeff | 599 | 10.0000 | 16.8580 |
| bond_coeff | 600 | 10.0000 | 20.0050 |
| bond_coeff | 601 | 10.0000 | 18.4150 |
| bond_coeff | 602 | 10.0000 | 11.6640 |
| bond_coeff | 603 | 10.0000 | 9.6490 |
| bond_coeff | 604 | 10.0000 | 6.4280 |
| bond_coeff | 605 | 10.0000 | 9.0640 |
| bond_coeff | 606 | 10.0000 | 10.2070 |
| bond_coeff | 607 | 10.0000 | 7.5300 |
| bond_coeff | 608 | 10.0000 | 5.1940 |
| bond_coeff | 609 | 10.0000 | 8.2350 |
| bond_coeff | 610 | 10.0000 | 8.3890 |
| bond_coeff | 611 | 10.0000 | 5.4470 |
| bond_coeff | 612 | 10.0000 | 5.9800 |
| bond_coeff | 613 | 10.0000 | 9.2640 |
| bond_coeff | 614 | 10.0000 | 9.1750 |
| bond_coeff | 615 | 10.0000 | 6.4450 |
| bond_coeff | 616 | 10.0000 | 8.4880 |
| bond_coeff | 617 | 10.0000 | 11.5400 |
| bond_coeff | 618 | 10.0000 | 10.8300 |
| bond_coeff | 619 | 10.0000 | 10.1280 |
| bond_coeff | 620 | 10.0000 | 13.1740 |
| bond_coeff | 621 | 10.0000 | 15.3130 |
| bond_coeff | 622 | 10.0000 | 13.8960 |
| bond_coeff | 623 | 10.0000 | 12.6020 |
| bond_coeff | 624 | 10.0000 | 11.6820 |
| bond_coeff | 625 | 10.0000 | 8.1820 |
| bond_coeff | 626 | 10.0000 | 7.9620 |
| bond_coeff | 627 | 10.0000 | 9.0780 |

| | | | |
|---|---|---|---|
| bond_coeff | 628 | 10.0000 | 9.8030 |
| bond_coeff | 629 | 10.0000 | 8.2990 |
| bond_coeff | 630 | 10.0000 | 5.2770 |
| bond_coeff | 631 | 10.0000 | 11.1550 |
| bond_coeff | 632 | 10.0000 | 12.3650 |
| bond_coeff | 633 | 10.0000 | 10.0170 |
| bond_coeff | 634 | 10.0000 | 6.9530 |
| bond_coeff | 635 | 10.0000 | 9.0310 |
| bond_coeff | 636 | 10.0000 | 9.9290 |
| bond_coeff | 637 | 10.0000 | 6.8810 |
| bond_coeff | 638 | 10.0000 | 6.3820 |
| bond_coeff | 639 | 10.0000 | 10.1670 |
| bond_coeff | 640 | 10.0000 | 9.5200 |
| bond_coeff | 641 | 10.0000 | 11.6020 |
| bond_coeff | 642 | 10.0000 | 12.0590 |
| bond_coeff | 643 | 10.0000 | 8.5830 |
| bond_coeff | 644 | 10.0000 | 7.6490 |
| bond_coeff | 645 | 10.0000 | 11.0460 |
| bond_coeff | 646 | 10.0000 | 10.6770 |
| bond_coeff | 647 | 10.0000 | 7.5090 |
| bond_coeff | 648 | 10.0000 | 8.4020 |
| bond_coeff | 649 | 10.0000 | 11.8590 |
| bond_coeff | 650 | 10.0000 | 11.0980 |
| bond_coeff | 651 | 10.0000 | 7.7460 |
| bond_coeff | 652 | 10.0000 | 9.8460 |
| bond_coeff | 653 | 10.0000 | 11.9410 |
| bond_coeff | 654 | 10.0000 | 6.7340 |
| bond_coeff | 655 | 10.0000 | 10.2650 |
| bond_coeff | 656 | 10.0000 | 11.7410 |
| bond_coeff | 657 | 10.0000 | 10.1310 |
| bond_coeff | 658 | 10.0000 | 11.3350 |
| bond_coeff | 659 | 10.0000 | 14.5410 |
| bond_coeff | 660 | 10.0000 | 15.0910 |
| bond_coeff | 661 | 10.0000 | 14.3640 |
| bond_coeff | 662 | 10.0000 | 16.7150 |
| bond_coeff | 663 | 10.0000 | 18.9760 |
| bond_coeff | 664 | 10.0000 | 19.3280 |
| bond_coeff | 665 | 10.0000 | 19.7440 |
| bond_coeff | 666 | 10.0000 | 22.4200 |
| bond_coeff | 667 | 10.0000 | 24.0660 |
| bond_coeff | 668 | 10.0000 | 5.7970 |
| bond_coeff | 669 | 10.0000 | 4.7080 |
| bond_coeff | 670 | 10.0000 | 8.4540 |
| bond_coeff | 671 | 10.0000 | 8.9390 |
| bond_coeff | 672 | 10.0000 | 7.3830 |
| bond_coeff | 673 | 10.0000 | 9.3220 |
| bond_coeff | 674 | 10.0000 | 12.1720 |
| bond_coeff | 675 | 10.0000 | 11.7400 |
| bond_coeff | 676 | 10.0000 | 10.8250 |
| bond_coeff | 677 | 10.0000 | 14.1980 |
| bond_coeff | 678 | 10.0000 | 15.9190 |

```
bond_coeff   679   10.0000  14.5530
bond_coeff   680   10.0000  15.7840
bond_coeff   681   10.0000  19.0010
bond_coeff   682   10.0000  22.3870
bond_coeff   683   10.0000  20.4620
bond_coeff   684   10.0000  18.5890
bond_coeff   685   10.0000  17.1720
bond_coeff   686   10.0000  16.2910
bond_coeff   687   10.0000  15.1620
bond_coeff   688   10.0000  13.7410
bond_coeff   689   10.0000  11.7410
bond_coeff   690   10.0000  11.1810
bond_coeff   691   10.0000  10.9580
bond_coeff   692   10.0000   9.4960
bond_coeff   693   10.0000   5.8490
bond_coeff   694   10.0000   7.2040
bond_coeff   695   10.0000  16.8220
bond_coeff   696   10.0000  16.2650
bond_coeff   697   10.0000  12.8570
bond_coeff   698   10.0000  11.7580
bond_coeff   699   10.0000  13.5710
bond_coeff   700   10.0000  13.3220
bond_coeff   701   10.0000   9.6480
bond_coeff   702   10.0000   9.6090
bond_coeff   703   10.0000  12.7900
bond_coeff   704   10.0000  13.5780
bond_coeff   705   10.0000  10.1470
bond_coeff   706   10.0000   9.7170
bond_coeff   707   10.0000  13.4520
bond_coeff   708   10.0000  14.6110
bond_coeff   709   10.0000  12.3360
bond_coeff   710   10.0000   8.7680
bond_coeff   711   10.0000  10.0520
bond_coeff   712   10.0000  11.5700
bond_coeff   713   10.0000   8.5120
bond_coeff   714   10.0000   6.2460
bond_coeff   715   10.0000   9.4360
bond_coeff   716   10.0000  10.6060
bond_coeff   717    0.0100  23.5970
bond_coeff   718    0.0100  21.3860
bond_coeff   719    0.0100  18.4650
bond_coeff   720    0.0100  19.9740
bond_coeff   721    0.0100  20.4190
bond_coeff   722    0.0100  16.9270
bond_coeff   723    0.0100  15.6920
bond_coeff   724    0.0100  18.6200
bond_coeff   725    0.0100  17.9370
bond_coeff   726    0.0100  14.2620
bond_coeff   727    0.0100  15.7740
bond_coeff   728    0.0100  18.6480
bond_coeff   729    0.0100  16.6210
```

| | | | |
|---|---|---|---|
| bond_coeff | 730 | 0.0100 | 15.5480 |
| bond_coeff | 731 | 0.0100 | 24.6300 |
| bond_coeff | 732 | 0.0100 | 21.9690 |
| bond_coeff | 733 | 0.0100 | 19.8430 |
| bond_coeff | 734 | 0.0100 | 20.9580 |
| bond_coeff | 735 | 0.0100 | 20.1540 |
| bond_coeff | 736 | 0.0100 | 16.6590 |
| bond_coeff | 737 | 0.0100 | 16.1770 |
| bond_coeff | 738 | 0.0100 | 18.2010 |
| bond_coeff | 739 | 0.0100 | 16.2550 |
| bond_coeff | 740 | 0.0100 | 12.9810 |
| bond_coeff | 741 | 0.0100 | 14.8150 |
| bond_coeff | 742 | 0.0100 | 16.0040 |
| bond_coeff | 743 | 0.0100 | 12.5040 |
| bond_coeff | 744 | 0.0100 | 12.1930 |
| bond_coeff | 745 | 0.0100 | 24.5430 |
| bond_coeff | 746 | 0.0100 | 23.5240 |
| bond_coeff | 747 | 0.0100 | 21.7200 |
| bond_coeff | 748 | 0.0100 | 22.9370 |
| bond_coeff | 749 | 0.0100 | 24.0230 |
| bond_coeff | 750 | 0.0100 | 23.5080 |
| bond_coeff | 751 | 0.0100 | 20.1400 |
| bond_coeff | 752 | 0.0100 | 19.1600 |
| bond_coeff | 753 | 0.0100 | 21.0680 |
| bond_coeff | 754 | 0.0100 | 19.2640 |
| bond_coeff | 755 | 0.0100 | 16.1430 |
| bond_coeff | 756 | 0.0100 | 20.0430 |
| bond_coeff | 757 | 0.0100 | 19.3360 |
| bond_coeff | 758 | 0.0100 | 16.3480 |
| bond_coeff | 759 | 0.0100 | 15.2720 |
| bond_coeff | 760 | 0.0100 | 15.8350 |
| bond_coeff | 761 | 0.0100 | 14.0810 |
| bond_coeff | 762 | 0.0100 | 10.9160 |
| bond_coeff | 763 | 0.0100 | 11.6300 |
| bond_coeff | 764 | 0.0100 | 12.2640 |
| bond_coeff | 765 | 0.0100 | 9.2230 |
| bond_coeff | 766 | 0.0100 | 7.2030 |
| bond_coeff | 767 | 0.0100 | 9.7060 |
| bond_coeff | 768 | 0.0100 | 8.5930 |
| bond_coeff | 769 | 0.0100 | 4.9330 |
| bond_coeff | 770 | 10.0000 | 16.6930 |
| bond_coeff | 771 | 10.0000 | 14.0970 |
| bond_coeff | 772 | 10.0000 | 8.5150 |
| bond_coeff | 773 | 10.0000 | 6.0790 |
| bond_coeff | 774 | 10.0000 | 19.8170 |
| bond_coeff | 775 | 10.0000 | 20.1490 |
| bond_coeff | 776 | 10.0000 | 21.9610 |
| bond_coeff | 777 | 10.0000 | 21.6880 |
| bond_coeff | 778 | 10.0000 | 16.3970 |
| bond_coeff | 779 | 10.0000 | 14.2300 |
| bond_coeff | 780 | 10.0000 | 3.9830 |

```
bond_coeff  781   10.0000  14.0580
bond_coeff  782   10.0000  17.8080
bond_coeff  783   10.0000  10.8120
bond_coeff  784   10.0000  22.8540
bond_coeff  785   10.0000  20.0120
bond_coeff  786   10.0000   8.1780
bond_coeff  787   10.0000  17.1920
bond_coeff  788    1.0000   6.7390
bond_coeff  789    1.0000   5.1680
bond_coeff  790    1.0000   8.8910
bond_coeff  791    1.0000  10.0550
bond_coeff  792    1.0000   8.1750
bond_coeff  793    1.0000   8.4020
bond_coeff  794    1.0000  11.7580
bond_coeff  795    1.0000  12.7210
bond_coeff  796    1.0000  11.9380
bond_coeff  797    1.0000  11.7320
bond_coeff  798    1.0000  14.7680
bond_coeff  799    1.0000  17.0700
bond_coeff  800    1.0000  14.5020
bond_coeff  801    1.0000  15.6940
bond_coeff  802    1.0000  13.8410
bond_coeff  803    1.0000  16.8950
bond_coeff  804    1.0000  16.6950
bond_coeff  805    1.0000  13.1830
bond_coeff  806    1.0000  13.6300
bond_coeff  807    1.0000  17.2580
bond_coeff  808    1.0000  16.3060
bond_coeff  809    1.0000  13.6220
bond_coeff  810    1.0000  14.6010
bond_coeff  811    1.0000  18.0930
bond_coeff  812    1.0000  18.8480
bond_coeff  813    1.0000  16.6270
```
